# Single-cell-level condition-related signal estimation with batch effect removal through neural discrete representation learning

**DOI:** 10.1101/2025.03.05.641743

**Authors:** Xiao Xiao

## Abstract

Advances in single-cell sequencing techniques and the growing volume of single-cell data have created unprecedented opportunities for uncovering the changes in gene expression patterns induced by perturbations or associated with diseases. However, batch effects and non-linearity in single-cell data make single-cell-level estimation challenging. To address these drawbacks, we developed NDreamer, an approach that combines neural discrete representation learning with counterfactual causal matching. NDreamer can be used to estimate the batch-effect-free and condition-related or perturbation-induced signal-preserved expression data from the raw expressions and then estimate single-cell-level perturbation-induced or condition-related signals. Benchmarked on datasets across platforms, organs, and species, NDreamer robustly outperformed previous single-cell-level perturbation effect estimation methods and batch effect denoising methods. Finally, we applied NDreamer to a large Alzheimer’s disease cohort and uncovered meaningful gene expression patterns between the dementia patients and health controls.

## Introduction

Single-cell RNA sequencing (scRNA-seq) has become a cornerstone technology for studying cellular heterogeneity and detecting differential gene expression patterns at the single-cell level. Studies have been conducted to sequence and identify expression changes induced by perturbations including genetic modifications^1^, chemical treatments^2^, and environmental stimuli^3^. Advanced technologies such as Perturb-seq^1, 4^ have further enabled us to study cellular responses to mutations in disease-related risk genes across diverse cell types.

In single-cell studies, one of the most important and challenging questions is to estimate how perturbation influences individual cells and how these signals vary across different cells. It is well known that cells with distinct intrinsic features, such as cell type, subtype, cell cycle stage, microenvironment, chromatin accessibility, and pre-existing states, respond differently to perturbations. Here, we define the perturbation-induced effect in each single cell as individual treatment effects (ITE) and its intrinsic features as effect modifiers in the framework of causal inference^5^. However, since we only observe the perturbation label and total gene expression of each cell. Without direct information on intrinsic features, it is difficult to disentangle intrinsic variability from perturbation-induced effects. Estimating ITE is further complicated by a key limitation of scRNA-seq: it captures a cell’s transcriptome under only one perturbation group, destroying the cell in the process and leaving its counterfactual state unknown. This necessitates the use of counterfactual causal inference techniques.

Early approaches relying on aggregating single-cell data to pseudo-bulk data to compare cells in the perturbation condition with cells in the control condition, where they averaged cells’ expressions for each sample according to their cell type annotations, often fail to capture fine-grained heterogeneity and can produce different results depending on the chosen aggregating resolutions. For example, CD4+ and CD8+ T cells respond differently to CD28 costimulation—CD4+ T cells sustain proliferation, whereas CD8+ T cells show a limited response^6^. Thus, aggregating all T cells could mask these subtype-specific responses, leading to insignificant results in the statistical analysis of T cell responses. Later methods tried to estimate single-cell-level ITE using single-cell digital twins produced by generative models^7, 8^ or counterfactual matching (or rigorously speaking, weighting) based on perturbation-independent factors^3, 9^. While these models have advanced the field of single-cell ITE estimation by identifying and disentangling perturbation-independent factors and isolating perturbation-related ITE responses from the expression matrix, they have several limitations. First, the unknown or subtle bias^10, 11^ in the expression profile generated by the generative model^7, 8^ can impact the ITE estimation and downstream analysis. Second, the inherent non-linearity of the single-cell data makes linear decompositions^12^ not suitable for analysis. Third, the evaluation of independence using correlation coefficients does not necessarily indicate independence, and utilizing the discriminator in adversarial training to establish independence may not rigorously lead to independence under various adversarial training schedules. Lastly, it is unknown if the distances in the perturbation-independent factors are distorted, which may harm downstream analysis that performs causal weighting or matching based on the distorted.

Additionally, in large single-cell studies with multiple batches, the batch effect can also bias the estimation of ITE but previous ITE estimation methods did not account for it. A large number of methods have been developed to remove batch effects^12, 13^. However, most of them provided a batch-effect-free embedding instead of denoised expressions, while over-correcting the perturbation-related signals along with the batch effect^12^.

Moreover, large-scale observational scRNA-seq studies comparing patient cohorts with healthy controls hold great promise for identifying disease (condition)-associated gene programs and biological pathways across different cell types, shedding light on disease mechanisms and potential therapeutic targets. However, making comparisons between groups and estimating single-cell-level condition-related signals (CRS) still remains challenging. Here, CRS can be viewed as ITE in the context of comparison between disease and control conditions, but there are only correlations and we can not make causal relationship conclusions based on CRS.

To overcome these challenges, we propose the NDreamer (Neural Discrete learning for decomposing condition-Related or perturbation-induced signals, Effect modifiers, And Measurement ERrors of complex forms in scRNA-seq) framework, to first decompose the measured gene expression into the true expression and batch effect, then estimate the batch-effect-free and condition-related or perturbation-induced signals for each individual cell. NDreamer can be applied to single-cell experiments, including multiple perturbation or condition labels with or without batches, and estimate the ITE/CRS for each cell and the conditional average treatment effect (CATE) conditional average condition-related signals (CACRS) for specific cell populations. We benchmark Ndreamer against existing single-cell-level condition-related signal estimation methods as well as batch effect-denoising methods and showed that our model outperformed all current methods. Applied to datasets across sequencing platforms, organs, diseases, and species, Ndreamer effectively identified cells’ heterogeneous responses to genetic edits, chemical treatments, and disease-associated gene expression patterns across different conditions while outperforming previous methods. We also applied NDreamer to the largest Alzheimer’s Disease scRNA-seq dataset to date and uncovered disease-related expression patterns across different cell types in the middle temporal gyrus (MTG) region.

## Results

### The unified modeling of gene expressions in single-cell sequencing studies

In single-cell studies, data are typically represented as two matrices: a cell-by-gene expression matrix and a cell-by-metadata annotation matrix. The metadata provides information on various factors, including condition labels and batches. With two or more categories, Conditions could be chemical perturbations, environmental stimuli, genetic modifications, and disease status. Batches, on the other hand, arise from several categorical factors such as donor ID, sequencing time, and data modality (Fig. 1a).

**Figure 1.**
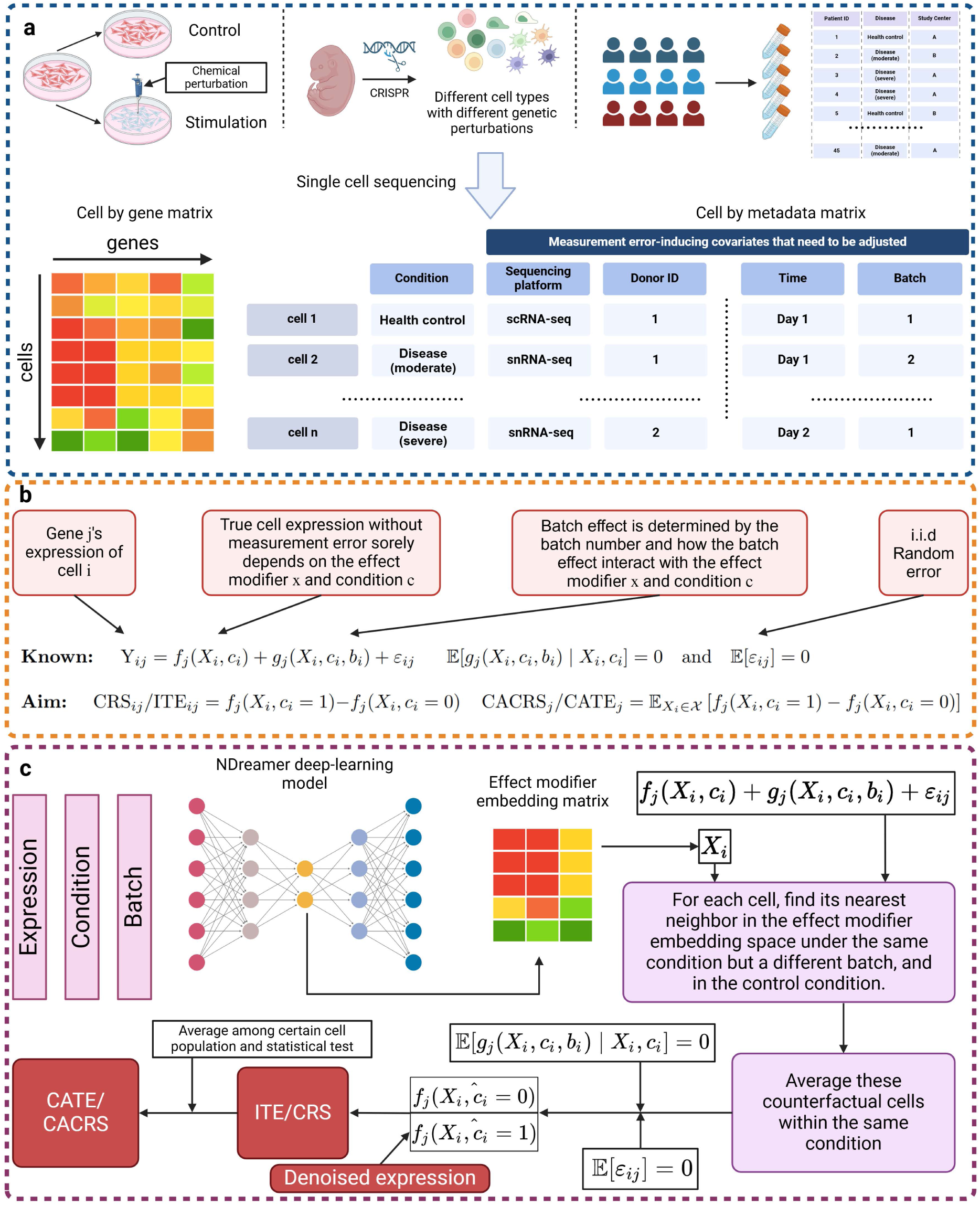
The general form of gene expressions in single-cell sequencing studies and the pipeline of NDreamer. **a.** Data structure in the single cell studies. We normally obtain a cell-by-gene expression matrix and a cell-by-metadata annotation matrix. The aim is to estimate the perturbation-induced or condition-related expression patterns. The perturbation or condition can be chemical perturbation, genetic perturbation, and disease status. **b.** Graphical illustration of the formula of the general form of gene expressions in single-cell sequencing studies. **c.** The pipeline of NDreamer in estimating batch-effect-free and condition-related or perturbation-induced signal-preserved expression profiles and ITE/CRS estimation.

Without loss of generality, we assume that the measured gene expression of a single cell in an experiment can be described as the combination of its true expression, batch effects (if present), and independent and identically distributed (i.i.d.) random noise (Fig. 1b). The true expression of a cell depends on the perturbation or condition label (e.g., perturbation and control) and its intrinsic characteristics, referred to as effect modifiers, such as cell type and cell state. The batch effect, arising from inevitable technical biases in measurement, is determined by the interaction between batch-specific attributes, the cell’s effect modifier features, and its experimental condition. Formally, the expression profile of the 𝑗-th gene in cell 𝑖, denoted as *Y*_*ij*_, can be modeled as:

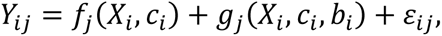

Where:

- 𝑋_*i*_ represents the effect modifier features of cell *i*,
- 𝑐_*i*_ denotes the condition (disease or perturbation v.s. control) label in which cell *i* resides,
- *f*_*j*_(*X*_*i*_, *c*_*i*_) is the true expression of the *j*-th gene in cell *i*,
- *b*_*i*_ indicates the batch to which cell *i* belongs,
- *g*_*j*_(*X*_*i*_, *c*_*i*_, *b*_*i*_) models the batch effect on the *j*-th gene in cell *i*, and
- 𝜀_*ij*_ represents i.i.d. random noise.

The functions *f*_*j*_ and *g*_*j*_ may be non-linear transformations, and this decomposition does not assume any linear combination of the terms mentioned above.

Since both the batch effect and the random noise are components of measurement error and noise, we assume 𝔼[*g*_*j*_(*X*_*i*_, *c*_*i*_, *b*_*i*_) ∣ *X*_*i*_, *c*_*i*_] = 0 and 𝔼[𝜀_*ij*_] = 0 . These assumptions are necessary: if the mean of the measurement error or noise were non-zero, it would be impossible to draw any reliable conclusion.

The first challenge is to unmix the condition-related responses and the condition-independent effect modifiers from the measured gene expression. To achieve this, we applied a neural network (see Methods) to extract the effect modifier that is independent of batch and condition while preserving the biological variation within each condition and batch.

Then, we estimated the denoised expression *f*_*j*_(*X*_*i*_, *c*_*i*_ = 1) for each cell by finding its nearest neighbor in each batch under the same condition in the effect modifier space and averaging them. It is hard to find the cell with exactly the same *X*_*i*_, so an alternative way is to find its nearest neighbor. And since 𝔼[*g*_*j*_(*X*_*i*_, *c*_*i*_, *b*_*i*_) ∣ *X*_*i*_, *c*_*i*_] = 0 and 𝔼[𝜀_*ij*_] = 0, the average can be considered as an estimate of *f*_*j*_(*X*_*i*_, *c*_*i*_ = 1). We then estimate its counterfactual cell’s expression in the control condition *f*_*j*_(*X*_*i*_, *c*_*i*_ = 0) by the same operation but for each batch under the control condition. By subtracting the counterfactual expression *f*_*j*_(*X*^, *c*_i_ = 0) from the estimated denoised expression, we get the ITE/CRS for cell *i*. Aggregate the ITE/CRS of certain cell populations and we get the CATE, and the corresponding nonparametric test can be applied to test its statistical significance (Fig. 1c). For convenience, we will only discuss the estimation of ITE since the estimation methods of ITE and CRS are the same under the proposed framework.

### Ndreamer deep learning model

Effectively extracting batch- and condition-independent effect modifiers while preserving within-batch and within-condition biological variation, without introducing distortions, remains a significant challenge. Without loss of generality, we assume that the effect modifier is independent of the batch and the condition^3^. Moreover, to make sure that the transform of the condition- and batch-independent effect modifier embedding is not an arbitrary transformation, we assume that the effect modifier space should contain the information of within-batch and within-condition between-cell variance where the within-batch and within-condition local and global distance structures should be retained in the effect modifier embeddings. To address this, we employed neural discrete representation learning^14, 15^, which generates categorical latent variables from the input expression profiles. Following discretization, a subsequent neural network transformation was applied to produce the final effect modifier embedding as a continuous vector. To constrain the embedding space and reduce distance distortions, we applied the variational inference techniques^16^ to the final effect modifier embedding and used a neural network decoder to reconstruct the expression (Fig. 2a).

**Figure 2.**
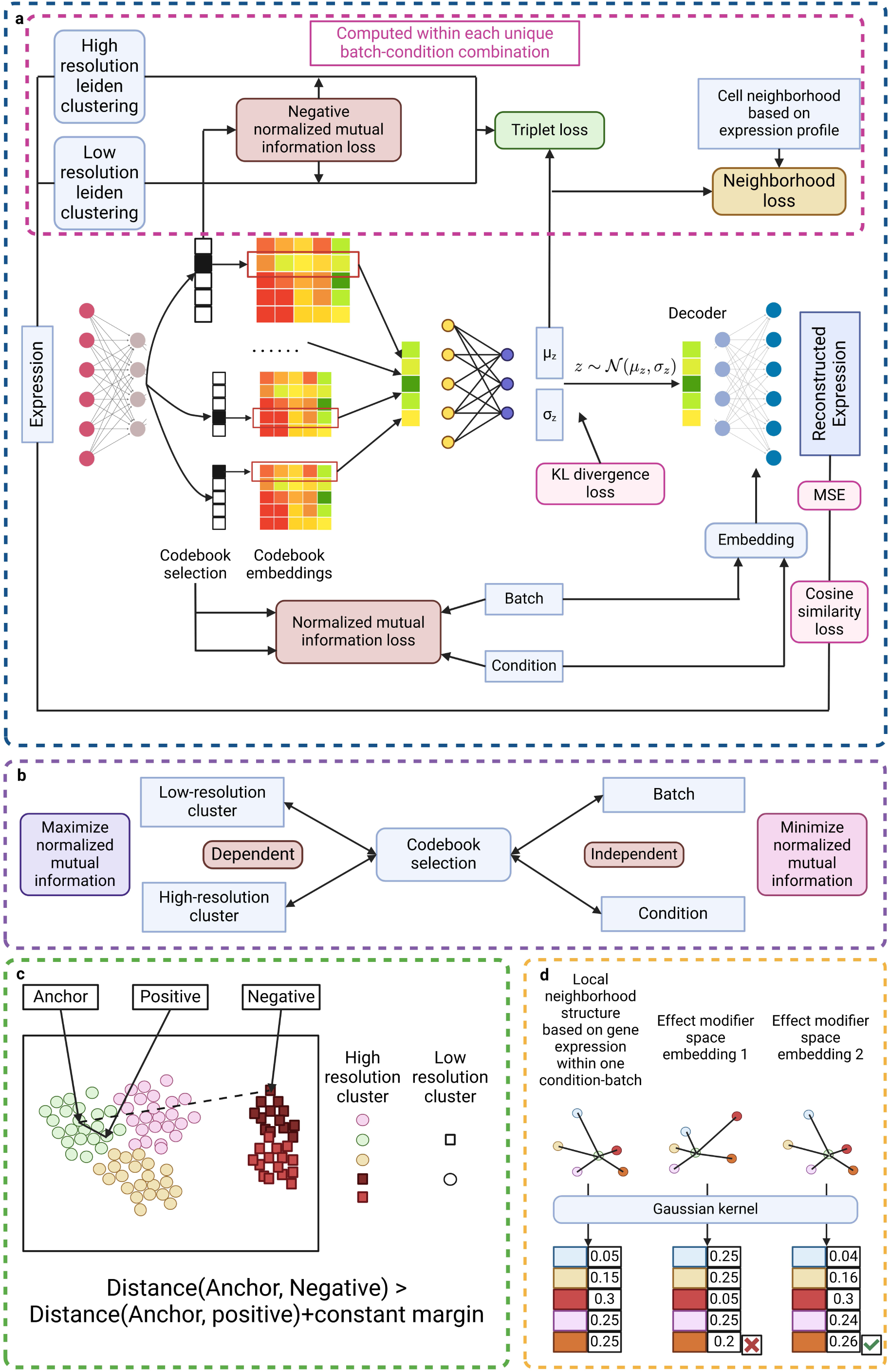
Overview of the NDreamer deep learning model for effect modifier extraction. **a.** The structure of the deep learning model. Input expression is first transformed to a set of categorical latent variables according to the neural discrete representation learning, then it is transformed to a continuous embedding, which is treated as the final effect modifier embedding, and then re-parametrized similar to VAE. Then a decoder is used to reconstruct the expression. For the categorical latent variables, we apply the independent and dependent loss using normalized mutual information. For the effect modifier embedding, we applied the triplet loss and the local neighborhood loss to preserve biological conservation. **b.** Graphical illustration of the loss based on the normalized mutual information. The categorical latent variables should be related to the unsupervised high- and low-resolution clusters in the raw expression space within each condition and batch, while being independent of the batch and the condition. **c.** Graphical illustration of the triplet loss function. Distance between cells within the same high- resolution unsupervised cluster should remain consistently smaller than the distance between cells from different low-resolution unsupervised clusters **d.** Graphical illustration of the local neighborhood loss. Local neighborhood structures calculated through the Gaussian kernel in the effect modifier space should mirror those in the raw expression space.

To ensure that the effect modifier embedding is independent of both batch and condition so that it can be used for matching and estimating ITE or CRS^3^, we first minimize the normalized mutual information between the categorical latent variable and the batch and condition (Fig. 2b). By decoupling the categorical latent variable from the batch and condition, the transformation of the categorical latent variable—the final effect modifier embedding— also becomes independent of these variables. Compared to the correlation-coefficient-based independence measurements as well as discriminator-based independence constraints^17^, the mutual information-based approach can more robustly capture complex independence patterns (Supplementary Notes 3).

After establishing independence through mutual information loss, we applied three loss functions to preserve within-batch and within-condition biological variation while preventing distortions. Firstly, the selection of the categorical latent variable was designed to depend on the unsupervised clustering patterns observed in the raw expression space within each condition and batch. Secondly, we employed a triplet loss to ensure that the distance between cells within the same high-resolution unsupervised cluster remained consistently smaller than the distance between cells from different low-resolution unsupervised clusters^18^. Together, these two loss functions safeguard the global biological structure. Beyond preserving global structure, we implemented a local neighborhood loss to maintain local structure, ensuring that the local neighborhood structures in the effect modifier space mirrored those in the raw expression space.

### NDreamer demonstrates robustly improved performance across experimental perturbation datasets

To systematically evaluate the performance of NDreamer with existing single-cell-level methods for perturbation-effect analysis, we conducted comprehensive benchmarking across diverse datasets across different sequencing platforms, species, and organs. Our comparison involves a set of currently state-of-the-art models, including CINEMA-OT^3^, Mixscape^9^, scCAPE^17^ (codes were downloaded from pip on 17^th^ November 2024), and scGen^7^ (implemented by pertpy^2^, codes were downloaded on 13^th^ December 2024). We apply the default hyperparameter settings provided by their GitHub tutorial page for these methods across all datasets. We treated the perturbation- and batch-free embeddings for previous methods as their estimated effect modifier embeddings. Our comparison is based on two categories of metrics:

1. Condition label-mixing performance in the effect modifier space. Given that the intrinsic characteristics of cells prior to any perturbation are independent of the perturbation label, the extracted effect modifier embeddings for cells in the two conditions (perturbation v.s. control), if plotted together, should be well-mixed. Please note that here we refer the perturbation and control as conditions here. Borrowing the ideas from the systematical batch-effect removal benchmarking papers^13, 19^, this can be evaluated by batch-mixing metrics where different perturbations are considered as different batches here. Specifically, we use the widely-used metrics of average silhouette width (ASW_batch)^19^, k-nearest-neighbor batch-effect test (kBET)^20^, batch local inverse Simpson’s index (bLISI)^21^, and True Positive Rate^22^ for batch-mixing evaluation.
2. Conservation of biological information for the effect modifier space. The extracted effect modifier embeddings should preserve the intrinsic features and within-perturbation variations of cells, which can be characterized by cell type annotation. Therefore, we employed the classical label conservation metrics to evaluate local neighborhoods (1 minus cell type LISI, cLISI)^21^, global cluster alignment (Adjusted Rand Index (ARI), geometric normalized mutual information (NMI)), and relative distances (cell-type ASW)^13^.

In addition to these metrics, we computed an overall score by linearly scaling the values of each metric to a range of 0 to 1 and then averaging the scaled values across all metrics. We used five publicly available datasets for the benchmarking: (1) The peripheral blood mononuclear cell (PBMC) dataset, including 16893 from IFN-beta-stimulated and control conditions^23^. (2) The rhinovirus infection (virus) dataset where we compared the 12255 cells between the mock condition and the exposure to 2% cigarette-smoke extract (CSE) perturbation^3^. (3) A small PBMC stimulation (PBMC1) dataset, which contains 5027 cells from the stimulated and control conditions^3^. (4) The ECCITE dataset, containing 20729 cells of 3 annotated cell cycle phases (G1, G2M, S) coming from 3 technique replicates measured using ECCITE-seq with 25 target genes generated from stimulated THP-1 cell line^9^. (5) The autism spectrum disorder (ASD) dataset containing brain cells with 35 kinds of neurodevelopmental delay (ND) risk gene mutation from 18 batches measured by Perturb-Seq^1^.

We begin with the PBMC dataset. As illustrated in Fig. 3a, 3b, NDreamer provides the uniform manifold approximation and projection^24^ (UMAP) visualization with the clearest cell type separation and the most effective mixing of cells from two different conditions, and achieves the highest overall score compared to other methods. Notably, NDreamer is the only method that successfully captures the variation between dendritic cells and other cell types. Among the other methods, CINEMA-OT achieves the second-best performance in mixing cells across conditions; however, it incorrectly splits a single cluster of CD4 T cells into two subgroups and fails to clearly distinguish between CD14+ Mono, FCGR3A+ Mono, and dendritic cells. The remaining three methods, while achieving relatively good cell type separation, fail to integrate cells from the two conditions effectively.

**Figure 3.**
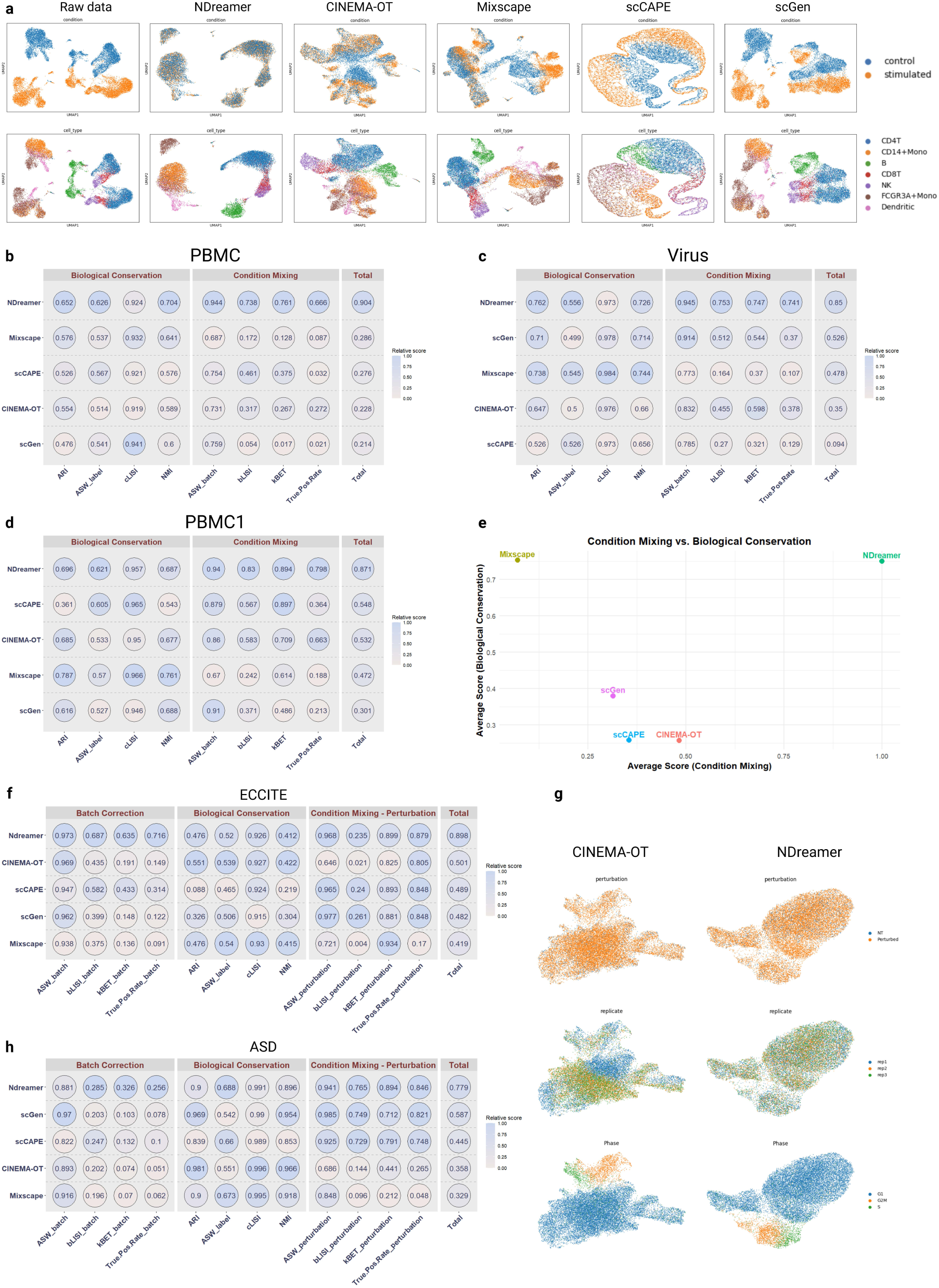
Benchmarking of NDreamer and other single-cell experimental perturbation analysis methods. **a.** UMAP visualization of the raw expression data, and the effect modifier embeddings of NDreamer, CINEMA-OT, Mixscape, scCAPE, and scGen (left to right). **b-d.** Quantitative evaluation of NDreamer and other methods on the PBMC (b), virus (c), and PBMC1 (d) datasets. **e.** Scatterplot of average biological conservation score against the average condition-mixing score for each method. **f.** Quantitative evaluation of NDreamer and other methods on the ECCITE dataset from the aspects of batch mixing, biological conservation, and condition mixing. **g.** UMAP of the effect modifier embedding of NDreamer and CINEMA-OT, colored by condition, batch, and cell cycle phase. **h.** Quantitative evaluation of NDreamer and other methods on the ASD dataset from the aspects of batch mixing, biological conservation, and condition mixing.

We then compared NDreamer to other methods using the virus and PBMC1 datasets. As shown in Fig. 3c–3d and Extended Figs. 1–2, NDreamer consistently outperformed the other methods with a 62% and 59% increase in the overall score. When averaging biological conservation and condition-mixing performance across all benchmarked methods on the three datasets, NDreamer emerged as the only method that achieved strong performance in both aspects. In contrast, Mixscape demonstrated limited condition-mixing capability, while the other three methods showed inadequate performance in preserving biological conservation.

However, in some single-cell experimental perturbation studies, multiple batches are present, and batch effects can significantly influence analysis results. This challenge is further amplified in genetic perturbation studies, where datasets often contain tens of genetic mutations. Previous methods, which only support comparisons between two conditions and lack batch-effect removal functionality, are less applicable in such scenarios. We illustrate this using the ECCITE and ASD datasets.

As shown in Extended Fig. 3a, the raw data exhibits a pronounced batch effect, with distinct separations between replication 1 and the other replications. To address this, we compared NDreamer (with batch-effect removal enabled) against four other methods in comparing genetically perturbed and unperturbed cells. In Fig. 3g, CINEMA-OT fails to mitigate batch effects, displaying a clear batch-related separation in the effect modifier space.

To systematically evaluate these methods when batch effect exists, we computed condition mixing, biological conservation, as well as batch-effect removal performance with the same metrics for condition mixing but applied to batches instead. As shown in Figs. 3f and 3h, NDreamer outperformed all other methods combining these three metrics, delivering embeddings with excellent condition-mixing and batch-mixing performance (Extended Fig. 3; Supplementary Fig. 1). In addition, we also benchmarked these methods by treating each genetic mutation as a condition and enable batch-effect-removal, if applicable, where we found that NDreamer still outperformed all other methods (Extended Fig. 4). The detailed results of the experimental perturbation study benchmarking is available in Supplementary Table 1.

**Figure 4.**
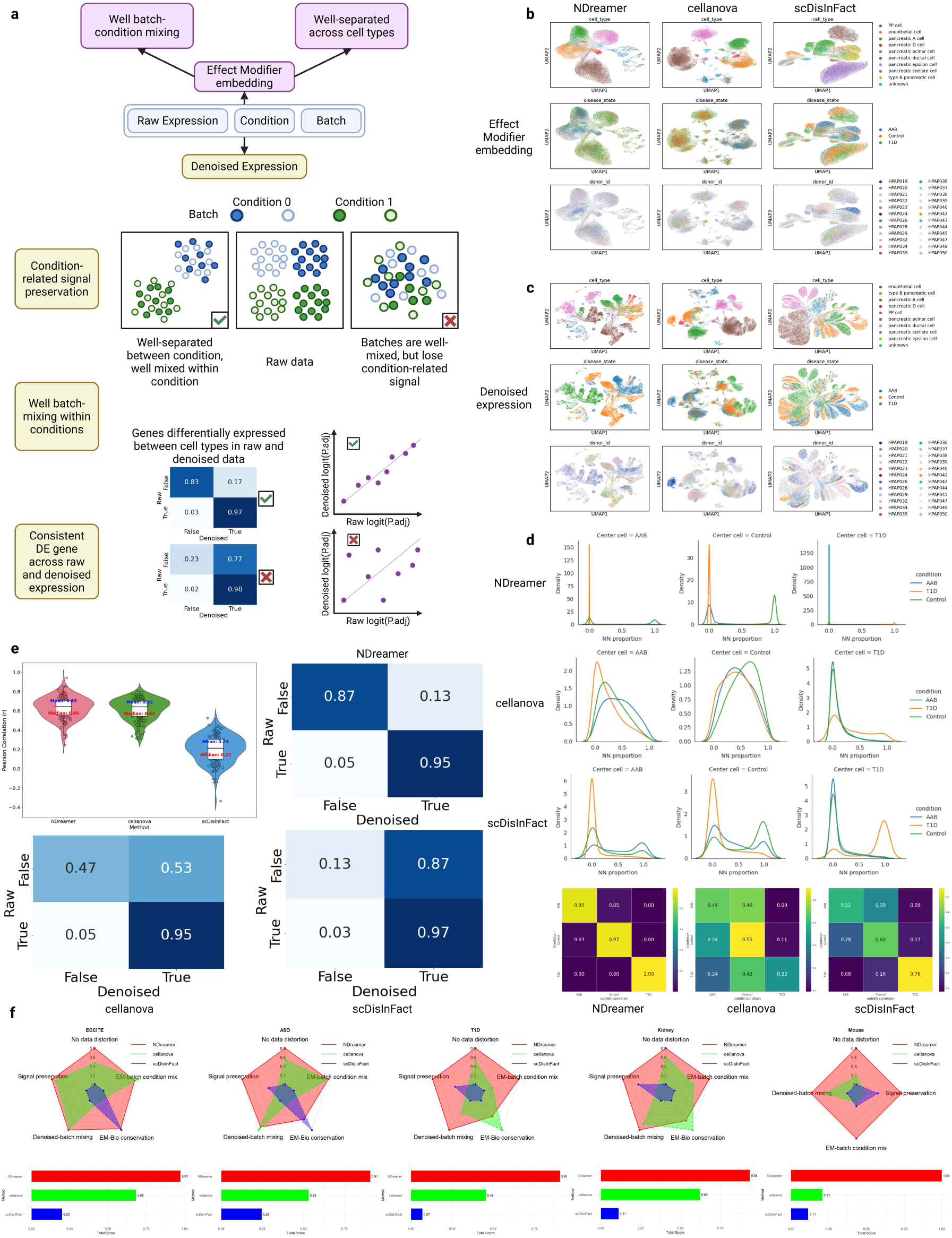
Benchmarking of NDreamer and other integration methods on producing batch-effect-free and condition-related-signal-preserved denoised expression. **a.** Illustration of the criteria of a good batch-effect-removal method. Firstly, the batch- and condition-free embedding used for downstream analysis should have good batch- and condition-mixing performance while retaining good biological conservation. Then, for the denoised expression, the batches should be well-mixed within one condition, cells in different conditions should be well-separated, and the variation between cell types should be retained. **b.** UMAP visualization of the effect modifier embeddings for NDreamer, CellANOVA, and scDisInFact colored by cell type, disease state, and donor id. **c.** UMAP visualization of the denoised expressions for NDreamer, CellANOVA, and scDisInFact colored by cell type, disease state, and donor id. **d.** Evaluation of condition-related-signal preservation. The distribution of the oobNN condition proportion for each condition (upper) and the neighboring cell condition versus central cell condition matrix visualization (lower) for NDreamer, CellANOVA, and scDisInFact. **e.** Evaluation of data distortion. The distributions of the PCC between raw and denoised expression’s each gene’s logit(adjusted p-value) obtained from DEG analysis for each cell type in each CBC across 3 methods (upper left). And the visualization of the normalized confusion matrix between the DEGs with BH-adjusted p-values less than 0.05 in the raw and denoised expression. **f.** Radar plot of method comparison from 5 perspectives: Data distortions in the denoised expression; Condition-related signal preservation for denoised expression (signal preservation); Batch-condition-mixing performance for the effect modifier embedding (EM-batch condition mix); Biological conservation for the effect modifier embedding (EM-Bio conservation); Batch-mixing performance within each condition for the denoised expression (Denoised-batch mixing).

### Systematical evaluation of NDreamer: Does it produce batch-effect-free and condition-related-signal-preserved expression?

In addition to extracting effect modifiers and estimating ITE in experimental perturbation studies, NDreamer can also be applied to observational studies to denoise true condition-related expressions from raw expressions influenced by batch effects and estimate CRS. To systematically evaluate its performance, we benchmarked NDreamer against current state-of-the-art models, including CellANOVA^12^ and scDisInFact^25^, using the default hyperparameter settings for these methods. To systematically evaluate which method can produce denoised expressions that are both free from batch effects and preserve condition-related signals, drawing inspiration from previous studies^12^, we conducted benchmarking based on five evaluation aspects (Fig. 4a):

1-2. **Batch-condition-mixing performance and biological conservation for the effect modifier embeddings.** NDreamer and previous methods all first identified a batch- and condition-free embedding for each cell, then generate a denoised expression based on these embeddings. As a result, the prerequisite of generating a reliable denoised expression, the embeddings must be well-mixed across conditions and batches, and preserve the within-batch or within-condition variations between cell types. To evaluate this, we applied the four batch-mixing metrics: ASW_batch, bLISI, kBET, and True positive rate for batch-condition-mixing evaluation and ARI, ASW_label, cLISI, NMI for biological conservation evaluation.

3. **Data distortion in denoised expression.** While denoised data inevitably differ from raw expression data, it is critical to preserve within-batch and within-condition variations between cell populations. Specifically, the differentially expressed genes (DEGs) between cell types should remain as consistent as possible with the original data. To quantitatively evaluate this, we computed the Benjamini-Hochberg (BH) adjusted p-values for each gene between cells in one cell type and all other cells within each condition-batch combination (CBC) for both raw and denoised data. The Pearson correlation coefficient (PCC) between the logits of adjusted p-values of raw and denoised data for each cell type and CBC was calculated, with the mean PCC across all CBCs serving as the first metric. Besides correlation, the preservation of the p-value’s scale is also important. To evaluate it, we categorized genes using a p-value cutoff of 0.05, with those having adjusted p-values below 0.05 considered differentially expressed genes (DEGs). Using the DEGs identified in the raw expression data as the ground truth and those from the denoised data as predictions, we calculated the prediction accuracy and F1 score as the second metric.

4. **Condition-related signal preservation for denoised expression.** In single-cell studies, cells from different conditions (e.g., disease versus healthy control) are expected to exhibit distinct differences, even when they belong to the same cell type or share similar intrinsic features. Capturing these cross-condition differences is crucial for understanding condition-specific biological variation. To assess whether these differences are preserved after integration, we first identify each cell’s 30 out-of-batch nearest neighbors (oobNNs) within the integrated embedding. Out-of-batch nearest neighbors are defined as cells from other batches (or samples) that are closest to a given cell in the integrated space. This approach ensures that comparisons are not confounded by batch-specific biases. For each cell, we then calculate the proportion of its oobNNs that belong to the same condition as the cell itself. If a cell represents a state with condition-specific differences, a higher proportion of its oobNNs is expected to come from the same condition. For quantitative comparison, we aggregate these proportions across cells and stratify them by condition, visualizing their density distributions. Effective integration is indicated by a higher proportion of neighbors belonging to the same condition as the central cell, reflecting the successful recovery of condition-specific signals. Conversely, failure to capture these signals results in uniform distributions, lacking enrichment for same-condition neighbors. To formalize this, we construct a neighboring cell condition versus central cell condition matrix. The value at the i-th row and j-th column represents the average proportion of oobNNs from the j-th condition for cells in the i-th condition. The trace of this matrix, normalized by the number of rows, serves as a performance measure: a value closer to 1 indicates that the matrix closely resembles an identity matrix, signifying better recovery of condition-specific signals.

5. **Batch-mixing performance within each condition for denoised expression.** After batch-effect removal, the cell’s expression should be only determined by its intrinsic features and the condition it belongs to. As a result, cells from different batches within the same condition should be well-mixed across batches. To evaluate this, we average the batch-mixing metrics ASW_batch, bLISI, kBET, and True positive rate that are applied to the batch across conditions.

In addition to these metrics, we computed an overall score by first averaging the metrics within the same aspect, then linearly scaling these values of each aspect to a range of 0 to 1, and then averaging the scaled values across all metrics.

Besides the above-mentioned ECCITE and ASD datasets, we also applied another 3 large datasets with obvious batch effect (Supplementary Fig. 2) to evaluate the batch-effect-removal and condition-related-signal-preservation performance: (1) The type 1 diabetes (T1D) dataset that contains 69,645 cells from 24 batches under 3 conditions (11 healthy individuals, 5 individuals with T1D, and 8 individuals with no clinical presentation of T1D but positive for β-cell autoantibodies (AAB)). (2) The human kidney multiomics atlas (kidney) dataset containing 282,610 cells from 36 donors and 47 batches with 24 ‘Control’ batches and 23 ‘Disease’ batches, where 29 of them are scRNA-seq data and 18 of them are snRNA-seq data. The analysis of this dataset includes multi-modal data integration. (3) The mouse radiation therapy (mouse) dataset containing 91,752 cells from the epithelial and lamina propria layers of the intestinal segments sequenced at 4 distinct times by two separate technical groups from mice under the control and radiation-treated condition. We will focus on illustrating the T1D dataset in the following paragraph.

We first visualized the batch- and condition-free embeddings produced by the three methods. As shown in Fig. 4b, NDreamer not only achieved the best condition and batch mixing performance compared to the other two methods but also demonstrated robust cell-type separation. While CellANOVA achieved a slightly higher score in biological conservation than NDreamer (Supplementary Table 2), it incorrectly split cell populations that should remain unified. For instance, the embeddings generated by CellANOVA divided pancreatic A cells and pancreatic epsilon cells into multiple distinct subgroups, despite the raw expression data indicating that these cell types are not segregated into separate clusters (Supplementary Fig. 2).

We then visualized the denoised expressions generated by the three methods using UMAP. As shown in Fig. 4c, NDreamer effectively separates cells from different conditions and retains the differences between different cell types, while CellANOVA only partially distinguishes between T1D and other conditions. In contrast, the denoised expression produced by scDisInFact still exhibits batch effects. Notably, NDreamer is the only method that successfully integrates cells from different batches within each condition, achieving well-mixed embeddings.

To quantitatively evaluate the denoised expression, we first focused on condition-related signal preservation. As shown in Fig. 4d, the oobNNs for NDreamer’s denoised expression are predominantly from the same condition, resulting in a neighboring cell condition versus central cell condition matrix that closely resembles an identity matrix, with a trace similarity of 0.973. In contrast, the other two methods exhibit less effective separation between conditions, with CellANOVA achieving a trace similarity of 0.442 and scDisInFact achieving a trace similarity of 0.624.

Next, we evaluated the extent of data distortion. We began by visualizing the distribution of PCCs between the adjusted p-values of genes in the raw and denoised expressions for each cell type and CBC. As shown in Fig. 4e, NDreamer achieved the highest mean and median PCC values, outperforming both CellANOVA and scDisInFact. Additionally, the normalized confusion matrix comparing raw (ground truth) and denoised (predicted) DEGs reveals that NDreamer introduced significantly less data distortion than the other two methods. Notably, while CellANOVA’s PCC values were slightly lower than those of NDreamer, its DEG prediction accuracy and F1 score (0.556 and 0.439, respectively) were substantially lower compared to NDreamer’s values (0.874 and 0.545). Combining the metrics of the 5 aspects together, NDreamer achieved a much higher overall score compared to other methods.

We further benchmarked NDreamer against CellANOVA and scDisInFact using additional datasets, including the ECCITE, ASD, kidney, and mouse datasets. Due to the absence of fine-grained cell-type annotations for the mouse dataset, biological conservation metrics were not evaluated for this dataset. Across all datasets, NDreamer consistently outperformed the other methods with the highest overall scores (Fig. 4f, Extended Figs. 5–8; Supplementary Figs. 3–6, Supplementary Table 2).

**Figure 5.**
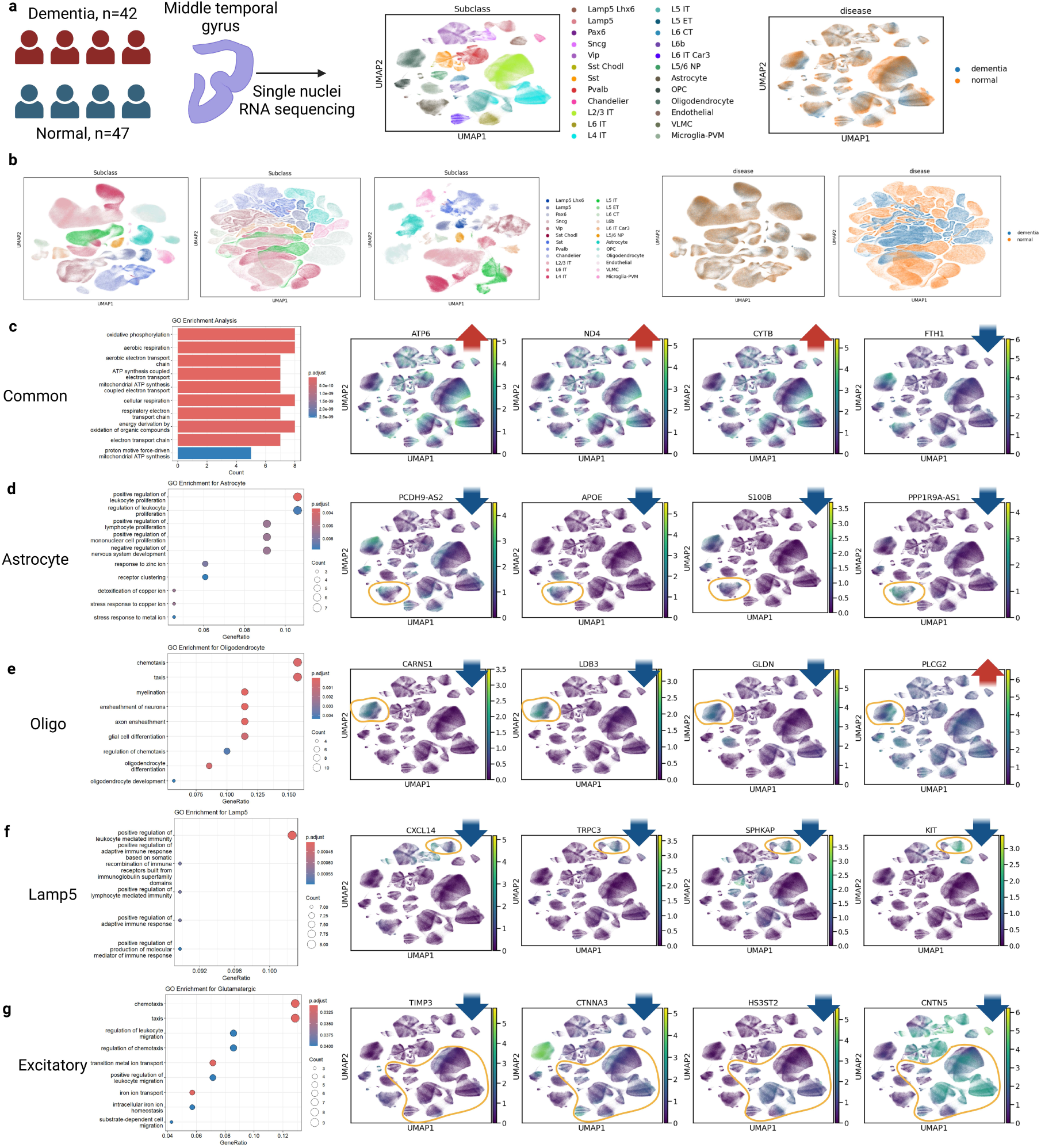
NDreamer uncovers differential gene expression patterns in the SEA-AD dataset. **a.** Overview of the SEA-AD MTG snRNA-seq dataset. There are 47 donors labeled as ’normal’ and 42 donors labeled as ’dementia’ in the ’disease’ annotation (left) with 24 kinds of cell type annotations (right). **b.** UMAP visualization of the effect modifier embedding, estimated denoised expression, and CRS colored by cell type (left), and the effect modifier embedding and estimated denoised expression colored by condition. **c.** The GO enrichment analysis results for the common top upregulated genes for all cell types (left) and the UMAP visualization of these genes on the UMAP of raw expressions. **d-g.** The GO enrichment analysis of the top down-regulated genes (left) and the UMAP visualization of the most significant DEGs except for the commonly up-regulated genes based the raw expression for (d) astrocytes, (e) oligodendrocytes, (f) Lamp5 inhibitory neurons, and (g) excitatory neurons.

### NDreamer uncovers differential gene expression patterns in a large Alzheimer’s disease cohort

In addition to benchmarking NDreamer against other methods, we applied it to the SEA-AD dataset^26^, the largest Alzheimer’s disease dataset to date. This dataset includes 1,378,211 single-nucleus RNA-sequenced (snRNA-seq) cells derived from the middle temporal gyrus (MTG) region of 47 donors labeled as ’normal’ and 42 donors labeled as ’dementia’ in the ’disease’ annotation (Fig. 5a). Identifying expression pattern shifts between cells in Alzheimer’s disease patients and healthy controls has long been a challenge. Our objective here is to estimate the CRS for each cell and make statistical tests for the CRSs in certain cell populations of interest. We began by visualizing the estimated effect modifier embedding, denoised expression, and CRSs of each cell using UMAP. As shown in Fig. 5b, the effect modifier embedding demonstrates excellent mixing of conditions and batches (Supplementary Fig. 7), while the denoised expression effectively separates cells from different conditions. Additionally, distinct clusters corresponding to different cell types (the ’Subclass’ annotation) are evident across the effect modifier embeddings, denoised expressions, and CRSs. These results indicate that NDreamer successfully extracts the condition- and batch-independent effect modifier, preserves condition-related signals and intrinsic cell feature variations, and effectively removes batch effects.

After estimating the CRS (Extended Fig. 9) and aggregating it by cell type to calculate the CACRS, we observed that most cell types share the same top-upregulated genes (Supplementary Table 3), including *ATP6*, *ND2*, *ND3*, *ND4*, *COX2*, and *CYTB*, while showing downregulation of the *FTH1* gene. These estimates align with the UMAP visualization of the raw expression data (Fig. 5c), where regions with higher dementia cell density exhibit altered expression levels of these genes compared to regions with higher normal cell density. Furthermore, our findings corroborate previous studies, where increased expression of mitochondrial genes^27, 28^ and disruption in iron metabolism are identified as the hallmarks of oxidative damage in AD brains^29^. To further investigate, we identified the intersection of the top 100 genes with the lowest p-values for the CACRS in each cell type and conducted a gene ontology (GO) enrichment analysis. This analysis revealed that these genes are strongly associated with cellular respiration and energy metabolism, consistent with prior research highlighting abnormalities in these processes as key features of AD pathology^30, 31^.

We then conducted a statistical analysis of the CRSs for astrocytes. In addition to the previously mentioned upregulated genes common across most cell types, the most significantly downregulated genes included *PCDH9-AS2*, *APOE*, *S100B*, and *PPP1R9A-AS1* (Fig. 5d), which have also been reported in previous studies^32–34^. Next, we examined the gene ontology (GO) terms enriched among the top 100 downregulated genes. These genes were primarily associated with the positive regulation of immune cell proliferation, as well as detoxification and stress responses to copper and other metal ions, which is also consistent with previous research^35, 36^.

Next, we analyzed the CRS and CACRS for oligodendrocytes (oligo). In addition to the commonly upregulated genes, we observed increased expression of *PLCG2* for oligo under the dementia condition, while *CARNS1*, *LDB3*, and *GLDN* were identified as the most significantly downregulated genes. These findings are consistent with previous studies^37–39^. Gene ontology (GO) enrichment analysis of the top 100 downregulated genes further highlights the vulnerability of oligodendrocytes and disruptions in myelin integrity in dementia conditions compared to normal conditions^37^. For the Lamp5 inhibitory neuron cell type, *CXCL14*, *TRPC3, SPHKAP*, and *KIT* genes are the most significantly down-regulated genes, which agree with previous studies^40^ and indicates the abnormality of the neuroinflammation^41^, calcium signaling^42^, and sphingosine kinase pathways^43^.

Lastly, we observed that most excitatory neurons shared the same top downregulated genes, including *TIMP3*, *CTNNA3*, *HS3ST2*, and *CNTN5*, which have been previously reported^44, 45^ and are known to be associated with amyloid precursor protein (APP) processing^46^ and tau pathology^47^. Gene ontology (GO) enrichment analysis of the top 100 downregulated genes further revealed that their differential expression would lead to dysfunction in pathways related to chemotaxis, inflammation, and ion transport.

### Exploration of batch effect, individual treatment effect, and sensitivity test

#### Exploration of Batch Effect

The batch effect for each cell can be directly estimated by subtracting the denoised expression from the raw expression, enabling visualization of the batch effect’s composition. As shown in Fig. 6a, batch effects were found to be cell-type-specific, with each batch exhibiting unique batch effects. This suggests that the true batch effect is a complex function of cells’ intrinsic features and batch-specific characteristics. To further explore this, we plotted the batch effect against factors contributing to batch composition, such as scRNA-seq versus snRNA-seq in the kidney dataset and sequencing time in the mouse dataset. Consistent with previous studies^12^, we found that the batch effect is closely associated with these factors, highlighting the multifaceted nature of batch effects.

**Figure 6.**
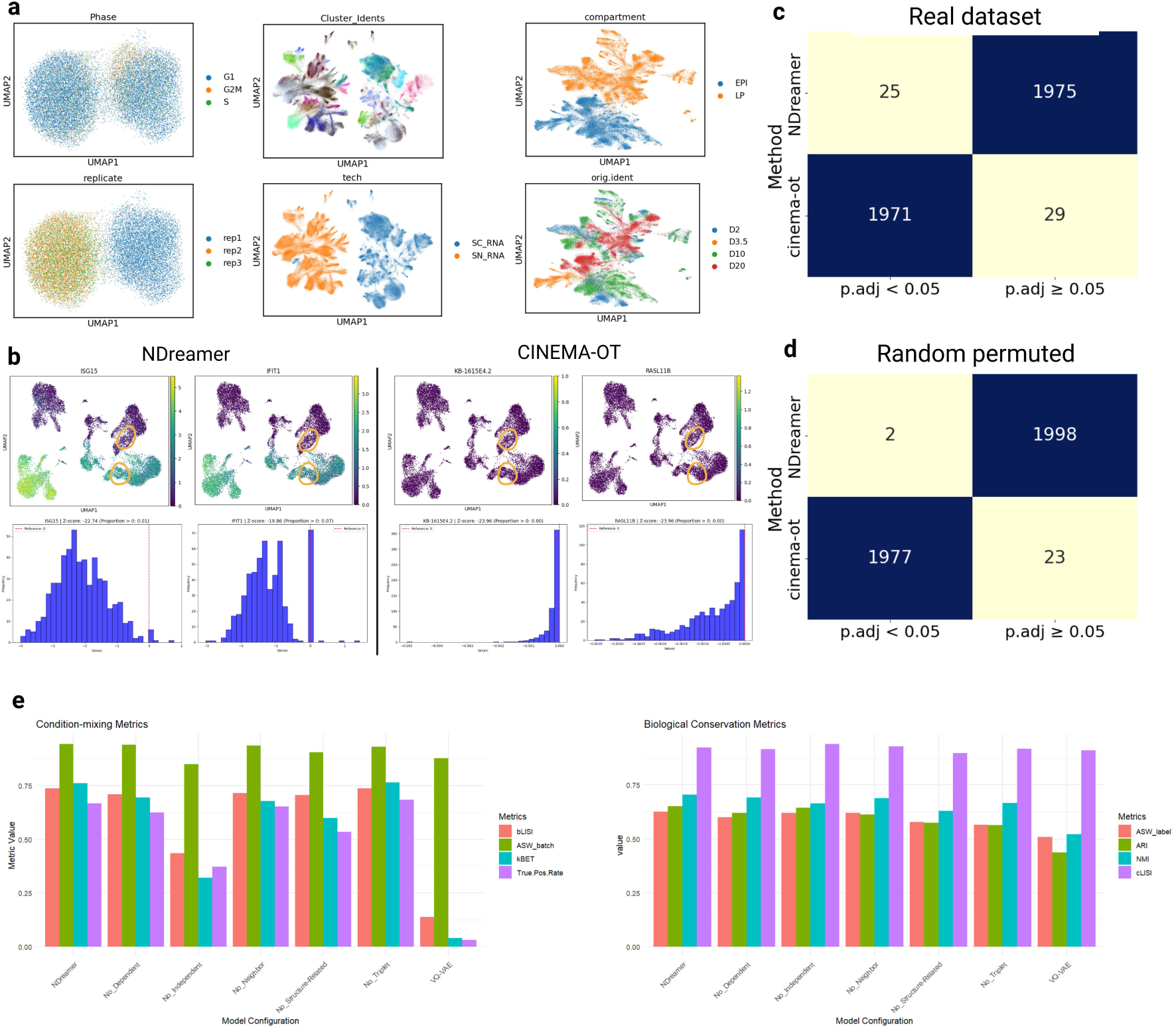
Exploration of batch effect, individual treatment effect, and sensitivity test. **a.** MAP visualizations of the estimated batch effect in the ECCITE dataset, colored by cell cycle phase and batch; the kidney dataset, colored by cell type and data modality; and the mouse datasets, colored by compartment and sequencing date. **b.** The number of DEGs with BH-adjusted p-values below 0.05, tested on CD8T cells in the PBMC dataset, for the NDreamer and CINEMA-OT methods. **c.** UMAP visualization of the top 2 most significant DEGs for the CD8T cells in the PBMC dataset estimated by NDreamer and CINEMA-OT. **d.** The number of DEGs with BH-adjusted p-values below 0.05, tested on CD8T cells in the randomly permuted PBMC dataset, for the NDreamer and CINEMA-OT methods. **e.** The ablation study of NDreamer on the PBMC dataset. The condition-mixing metrics and the biological conservation metrics are used. We include the models where we remove the dependent loss (No_Dependent), the local neighborhood loss (No_Neighbor), the triplet loss (No_Triplet), all the three mentioned biological conservation-related loss (No_Structure-Related), the independent loss (No_Independent), and all the previous mentioned loss (VQ-VAE).

#### Testing for Bias

A critical requirement for the estimated ITE is that it should be unbiased. To test this, we compared the ITEs estimated by NDreamer and CINEMA-OT, calculating BH-adjusted p-values for the ITEs of CD8T cells using a nonparametric method (see Methods). As shown in Fig. 6b, NDreamer identified only 25 genes with adjusted p-values less than 0.05, whereas CINEMA-OT reported 1971 out of 2000 highly variable genes as significant. This implausibly high number likely reflects bias in CINEMA-OT’s ITE estimation. We then visualized the two most significant genes with the lowest adjusted p-values (Fig. 6c) by plotting the UMAP colored by the expression level of these genes and the distribution of ITEs for these genes in CD8T cells. Genes identified by NDreamer exhibited clear differences between the two conditions, while those identified by CINEMA-OT appeared to lack meaningful significance, likely arising from bias. To further investigate, we randomly permuted condition labels for each cell and re-ran both methods. In this scenario, where significant p-values should be rare and driven only by random chance, NDreamer identified only two genes with adjusted p-values below 0.05. In contrast, CINEMA-OT still reported 1977 genes as significant (Fig. 6d). This finding strongly suggests that the ITEs estimated by CINEMA-OT are biased.

#### Nearest Neighbor Matching Is All You Need

Nearest neighbor matching, optimal transport matching, and k-nearest neighbor weighting are three widely used methods for covariate adjustment in estimating ITEs in causal inference. However, optimal transport and k-nearest neighbor weighting methods may introduce additional bias due to the sparsity and high proportion of zeros in single-cell data. Calculating counterfactual cells using (weighted) averages of multiple cells can result in a loss of sparsity in the counterfactual cells, potentially introducing bias when subtracting the counterfactual expression from the reference cell expression. As a result, theoretically, any weighting methods, if not one-to-one matching, would introduce the bias towards the negative direction in the ITE estimation, as shown in the ITEs of CINEMA-OT where most ITEs are negative. A detailed proof of why weighting-based methods are biased is provided in Supplementary Notes 4. In addition, since cells from different conditions and batches are well-mixed in the NDreamer-produced effect modifier space, there is no need to use optimal transport to adjust for the imperfect condition mixing performance as in previous methods.

#### Ablation study

To identify the key components of NDreamer’s deep learning model, we conducted an ablation study using the PBMC dataset. As shown in Fig. 6e, each loss function contributes to either condition-mixing performance or biological conservation. Specifically, removing the independent loss function (No_Independent) significantly decreases condition- mixing performance, demonstrating that the mutual information-based independent loss effectively identifies condition-independent effect modifiers. Similarly, eliminating the triplet loss (No_Triplet), local neighborhood loss (No_Neighbor), dependent loss (No_Dependent), or all three structure-related loss functions (No_Structure-Related) results in a notable reduction in biological conservation metrics. This highlights the critical role of these losses in preserving local and global biological structure. Moreover, comparing NDreamer to the original VQ-VAE model reveals a dramatic improvement across all metrics. This demonstrates that the inclusion of these loss functions successfully adapts NDreamer for causal inference in single-cell data.

## Discussion

Advances in single-cell sequencing techniques and the growing volume of single-cell data have created unprecedented opportunities for uncovering statistically significant gene expression patterns induced by perturbations or associated with specific disease conditions. However, existing analytical approaches face significant limitations. Many methods for estimating experimental effects at the single-cell level are either biased or ineffective at extracting condition-independent effect modifiers while preserving biological variation. Moreover, they struggle to scale for large experimental studies involving multiple batches with the batch effect. Similarly, current batch-effect removal methods often overlook the non-linear characteristics of single-cell data, fail to achieve adequate batch- and condition-mixing performance, or remove the condition-related signals while correcting for batch.

To address these challenges, we developed NDreamer, a robust framework designed to denoise expression profiles while preserving condition-related signals and effectively estimating individual treatment effects at the single-cell level. Through the combination of loss functions that ensure batch- and condition-independent while preserving global and local biological structure, NDreamer outperformed existing methods across all datasets. Additionally, with optimized tensor computation designs for the mutual information loss and the triplet loss calculations, NDreamer achieves exceptional computational efficiency, finishing model training on the PBMC dataset (2 conditions, 16,893 cells) in under one minute and on the large SEA-AD dataset (1,378,211 cells, 89 batches) in 3,837 seconds (Supplementary Notes 2) with GPU memory less than 8G when the batch size is set to 2048. The GPU memory only increases linearly with batch size and the computation time linearly increases with the number of batches, the number of conditions, and the number of cells in the dataset.

However, it is essential to carefully examine your data before applying NDreamer. To make the causal inference, NDreamer extracted the effect modifier embedding that is independent of both condition and batch. In some cases, however, the true effect modifier may not be entirely independent of these factors. For instance, in the ASD dataset, all cells of the “Vascular” cell type are unperturbed, making the condition inherently linked to the intrinsic features of the cell. Similarly, in the kidney dataset, certain cell types are common in the scRNA-seq modality but are rare in the snRNA-seq modality. Without additional information beyond the expression matrix, condition annotations, and batch annotations, these dependencies cannot be fully resolved based on information theory principles. Although the three loss functions in NDreamer are designed to retain biological conservation, they may still encounter limitations in such scenarios. The model can effectively extract effect modifiers while preserving biological conservation to a large extent, but the biological conservation metrics may experience a slight decrease. A practical suggestion in these situations is to exclude cells that inherently carry batch or condition information. For example, in the ASD dataset, if a cell is identified as belonging to the “Vascular” cell type, it can be excluded since it directly indicates the negative control condition. This preprocessing step can help mitigate potential confounding and enhance the reliability of downstream analyses.

Besides, this is the first study to integrate neural discrete representation learning with mutual information loss to establish independence, demonstrating the practical applicability of mutual information loss beyond its theoretical justification. Unlike previous methods that rely on adversarial training and discriminators to enforce independence, our approach is significantly more efficient, especially when the encoder is large (e.g. Transformers), as it reduces the need for multiple backpropagation steps required for discriminator training. Additionally, neural discrete representation learning offers distinct advantages over conventional VAEs, particularly when auxiliary discrete-related objective functions, such as to keep cluster consistency between effect modifier space and raw expression space, are involved. Finally, we established that one-to-one matching is the best method in experimental perturbation studies, otherwise, weighting may introduce bias towards the negative direction for the estimated ITE due to the sparsity of single-cell data. Although there is no good theoretical guarantee for the nearest neighbor matching, it is still currently the best causal inference method.

With its outstanding performance, efficient computation time, and the ability to solve the basic question in single-cell studies: what are the differences for cells with different intrinsic features between different conditions, we anticipate that, NDreamer will be widely adopted in single-cell studies to explore biological knowledge and discover key factors in diseases.

## Methods

### Data preprocessing

We processed the scRNA-seq data using the established Scanpy pipeline. Raw count matrices were first imported into Scanpy AnnData objects. To ensure data quality, we filtered out low-quality cells with fewer than 300 detected genes and excluded genes expressed in fewer than five cells. This step mitigated the risk of misleading alignment between cells with significant dropout events and those exhibiting inherently low transcriptional activity. Next, we normalized the library size of each cell to 10,000 reads by scaling gene counts based on total counts per cell and adjusting by a factor of 10,000. This normalization was followed by a log transformation, log(1 + 𝑥), stabilizing variance across the dataset. We then identified the top 2,000 highly variable genes. This preprocessing protocol was applied consistently across all benchmark datasets for NDreamer and other benchmarked methods unless other preprocessing (e.g. scaling) is mentioned in other methods (see the benchmarking section). For our method, we also performed unsupervised clustering within each batch using the Leiden algorithm, employing both a high-resolution parameter 𝑟_ℎ_, and a low-resolution parameter 𝑟_𝑙_.

### Estimation of the effect modifier embedding

#### Encoder with neural discrete representation learning and variational inference

For cell *i* with expression *Y*_*i*_, batch *b*_*i*_, and condition *c*_*i*_, we utilize a set of *n*_*c*_ codebooks, each containing *n*_e_ embeddings of dimensionality d_*c*_ , to discretize the input *Y*_*i*_ into a latent representation. First, for each codebook, *Y*_*i*_ is transformed by a codebook-specific neural network into an intermediate latent space *Z*_*i*_ ∈ ℝ^d*c*^. Each embedding in the codebook is then probabilistically selected based on the softmax of the negative Euclidean distance between *Z*_*i*_ and the embeddings. The probability of selecting the *j*-th embedding in a given codebook is computed as:

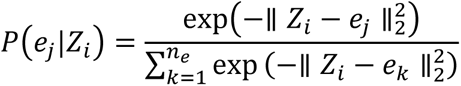

To make the random embedding selection operation differentiable, we employ the Gumbel-Softmax trick^48^, enabling backpropagation through the probabilistic codebook selection.

Putting together the results from each codebook, we generate a codebook choice matrix 𝑃 of shape *n*_*c*_ × *n*_e_ , recording which embedding *Z*_*i*_ is mapped for each codebook. After the embeddings are selected, we concatenate the chosen embeddings from each codebook to form a new latent space *Z*_*i*_′ ∈ ℝ^*nc*d*c*^, which serves as the discrete representation of *Y*_*i*_.

To ensure robust alignment between *Z*_*i*_ and the selected code e^∗^ and make the update of embeddings differentiable, a commitment loss with two components is employed similar to the VQ-VAE^14^. The first term penalizes the difference between *Z*_*i*_ and e^∗^, while the second term enforces the embeddings to remain close to *Z*_*i*_, preventing instability in the codebook updates. The full commitment loss is expressed as:

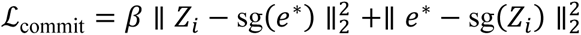

where sg(⋅) is the stop-gradient operator and 𝛽 (default to 0.25) controls the balance between the two terms.

To improve numerical continuity and reduce distortion in the estimated effect modifier embedding, we applied the variational inference technique similar to the standard Variational Autoencoder (VAE) framework. Based on the new latent space *Z*_*i*_′, we apply a multi-layer perceptron (MLP) to further transform *Z*_*i*_′ into two outputs: a mean vector 𝜇_*i*_ ∈ ℝ^d𝑧^ and a variance vector 𝜎_*i*_ ∈ ℝ^d𝑧^, where d_𝑧_ refers to the number of dimensions of the effect modifier embedding. The variance vector 𝜎_*i*_ is then exponentially transformed to ensure positivity. And the mean vector 𝜇_*i*_ is treated as the effect modifier embedding *X*_*i*_. Since *Z*_*i*_′ is independent of the condition and batch as constrained by the mutual information loss mentioned later, *X*_*i*_, as a transformation of *Z*_*i*_′, is also independent of the condition and batch. At last, we resample the latent effect modifier representation *X*_*i*_′ by:

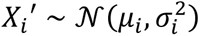

where reparameterization is used to enable backpropagation through the sampling operation, expressed as:

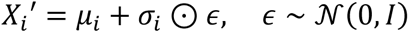

Following the Evidence Lower Bound (ELBO) loss in the training of traditional VAE^16^, the Kullback-Leibler (KL) divergence loss is introduced. The KL-divergence between the approximate posterior 𝑞(*X*_*i*_|*Y*_*i*_) ∼ 𝒩(𝜇_*i*_, 𝜎*_i_*^2^) and the prior 𝑝(*X*_*i*_) ∼ 𝒩(0, 𝐼) is given by:

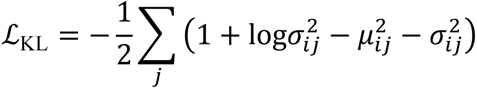

where *j* indexes the dimensions of the latent effect modifier embedding space. The resulting resampled effect modifier embedding *X*_*i*_′ serves as the final encoder output derived from the input *Y*_*i*_, integrating the benefits of both probabilistic embedding discretization and variational inference.

In addition, we found that some embeddings in the codebook are almost never used in model training, which reduces the model’s efficiency. To solve this problem, we reset the embeddings that are never selected every 30 steps when training, where we randomly replace the unused embeddings with the intermediate latent space embedding *Z*_*i*_ produced by the randomly chosen real data.

### Decoder with conditional generation

We then reconstruct the gene expression to let the model learn from the data using the resampled effect modifier embedding *X*_*i*_′ along with the batch embedding and the condition embedding. Written in formula:

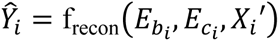

where 𝐸_*bi*_ and 𝐸_*ci*_ are the condition and batch embedding, and f_recon_ refers to the decoder neural network.

As part of the VAE ELBO, the reconstruction loss is defined as the mean squared error (MSE) between the measured expression *Y*_*i*_ and the reconstructed expression *Y*^_*i*_:

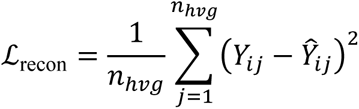

where *j* indexes the dimensions of the expression vector, and *n*_ℎ𝑣*g*_ is the total number of highly variable genes input to the model.

To further emphasize the interaction between the effect modifier *X*_*i*_, batch embedding 𝐸_*bi*_ , and condition embedding 𝐸_*ci*_ , we reinforce the reconstruction loss with an additional cosine distance loss.

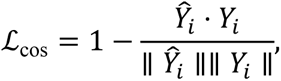

where · denotes the dot product, and ‖·‖ is the vector norm.

### Normalized mutual information loss

To enforce independence between the choice of the embeddings in the codebooks and the batch assignments as well as the condition assignments while preserving meaningful biological cluster information, we introduce the normalized mutual information loss. The choice of the embeddings in the codebooks is inherently categorical, allowing us to compute entropy and mutual information directly from the probability distributions. We start by introducing the calculation of mutual information between the choice of the embeddings in the codebooks and the batch.

Consider 𝐵 cells in a mini-batch (note that the mini-batch in deep learning training is different from the batch in the scRNA-seq study), let the batch assignment vector *b* ∈ ℝ^𝐵^ represent the experimental batch assignments. The assignment vector is first one-hot encoded into a matrix 𝔹 ∈ ℝ^𝐵×*nb*^ , where *n*_*b*_ is the number of unique batch categories. For one codebook, the joint probability 𝑃_*j*𝑜*in*𝑡_ ∈ ℝ^*nb*×*n*c^ , indicating the probability of a cell is simultaneously assigned to one of the *n*_*b*_ batches and one of the *n*_e_ embeddings in this codebook, can be then calculated by:

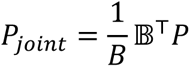

where 𝑃 ∈ ℝ^𝐵×*n*c^ is the probability matrix of codebook embedding selection for this codebook. This matrix-form operation enables acceleration by parallel computing.

The marginal probabilities of the batch and embedding assignment 𝑃(*b*) and 𝑃(c) are obtained by summing over the respective dimensions of 𝑃_*j*𝑜*in*𝑡_:

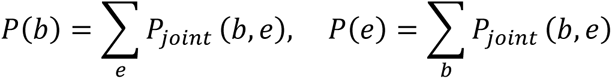

The marginal entropy of the batch and embedding assignment can be then calculated by:

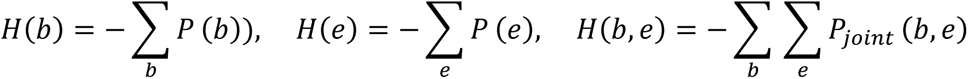

The mutual information is then calculated by:

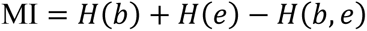

To stabilize the training and avoid nan, we use the arithmetic normalized mutual information (ANMI) as the loss function.

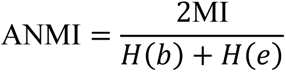

We then compute the ANMI of other codebook assignments with the batch and average them as the mutual information between the codebook assignments and the batch ℐ_*b*_ . By minimizing the mutual information, the loss encourages the latent variables to remain independent of the batch assignments.

Similarly, we calculate and minimize the ANMI between the codebook assignments and the condition assignment ℐ_*c*_. In addition, within each unique batch-condition combination, we maximize the ANMI between the codebook assignments and the unsupervised cluster label with low and high cluster resolution ℐ_𝑙𝑟_ and ℐ_𝑙ℎ_ to retain the global structure as in the original gene expression space.

### Triplet loss

The triplet loss is designed to preserve the relative distances in the effect modifier embedding, ensuring that the structure of the embedding space is not distorted^18^. The key idea is that cells belonging to the same small high-resolution cluster in the original space should be embedded closer to the anchor cell than cells from different low-resolution clusters. For a given cell *i*, the triplet consists of the anchor cell *i*, a positive cell *i*^+^ with effect modifier embedding *X*_*i*_+ from the same high-resolution cluster, and a negative cell *i*^−^ with effect modifier embedding *X*_*i*_− from a different low-resolution cluster. The triplet loss for a single triplet is defined as:

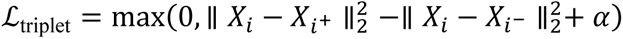

where *X*_*i*_ is the embedding of the anchor cell, *X*_*i*_+ and *X*_*i*_− are the embeddings of the positive and negative cells, respectively, ‖·‖_2_^2^ denotes the squared Euclidean distance, and 𝛼 > 0 is a margin that enforces separation between the anchor-negative and anchor-positive distances. For each cell *i*, *n*_𝑡_ such triplets are sampled, and the triplet losses are summed to compute the total triplet loss for the cell. This approach ensures that the effect modifier embeddings reflect meaningful relationships between cells while maintaining their relative distances in the embedding space.

### Local neighborhood loss

Besides using triplet loss and ANMI between codebook embedding selection and cluster information to preserve global biological conservation, we also designed the local neighborhood loss for retaining the local neighborhood structure in the effect modifier space within each unique batch-condition combination. For a reference cell *i*, we first identify its 𝑘_neighbor_ nearest neighbor in the original expression space and the effect modifier space, resulting in a cell subset 𝑚 (𝑘_neighbor_ ≤ |𝑚| ≤ 2𝑘_neighbor_). We then use the Gaussian kernel to calculate the relative distance between the reference cell and the 𝑚 cells in both spaces:

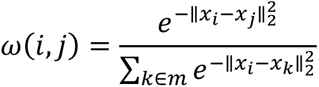

where 𝑥_*i*_ refers to the gene expression vector or the effect modifier embedding in cell *i*. This calculation results in the relative distance in the gene expression space and effect modifier space 𝛺_*g*_ and 𝛺_e_. The local neighborhood loss ℒ_neighbor_ is calculated as the mean squared error between 𝛺_*g*_ and 𝛺_e_ to encourage consistency between the neighborhood structure in the gene expression space and the effect modifier space.

### Implementation details

The total loss is calculated as:

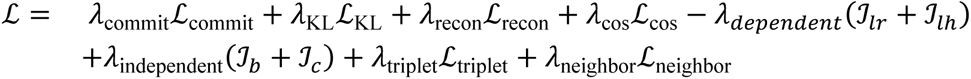

where each loss is multiplied with an empirical scaler similar to previous studies to balance the scale between different kinds of losses and optimize the model performance (Supplementary Notes 1-2).

We employed the Adam optimizer for training, configured with a learning rate of 0.001, 𝛽_1_ = 0.9, 𝛽_2_ = 0.999, and 𝜖 = 10^−8^. Here, 𝛽_1_ controls the exponential decay rate for the first moment estimates (momentum), 𝛽_2_ governs the decay rate for the second moment estimates (variance), and 𝜖 is a small constant added for numerical stability. To ensure consistent representation across training batches, each mini-batch contains 1024 cells with equal contributions from each unique batch-condition combination.

For the benchmarking part, metrics are calculated using the scib python package and the ’oobNN’ function in CellANOVA.

### ITE/CRS estimation

Since the computation of ITE and CRS, as well as CACRS and CATE are the same, we will only discuss about the calculation of ITE and CATE below. To estimate the ITE for each cell, we mitigate confounding effects arising from variations in intrinsic cell features by searching for the nearest neighbors in the effect modifier space in other batches under the same condition and the reference condition. The nearest neighbors are searched based on the Euclidean distance. These nearest cells are then averaged across batches within the same condition to obtain a batch-effect-free representation of the cell’s expression.

Let ℬ_treatment_ and ℬ_control_ represent the sets of batches that have cells in the treatment condition and the control condition, respectively. For each batch *b* ∈ ℬ_treatment_, the nearest cell *j*_*b*_ to cell *i* is defined as:

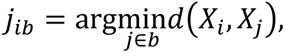

where d(*X*_*i*_, *X*_*j*_) is the Euclidean distance between *X*_*i*_ and *X*_*j*_. The batch-effect-free denoised expression under the treatment condition, *Y*_treatment_, is computed as:

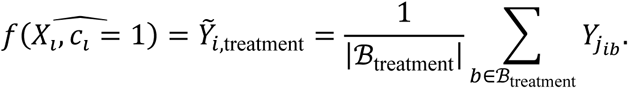

Similarly, the batch-effect-free denoised expression under the control condition is:

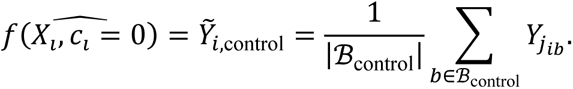

The individual treatment effect for cell *i* is then given by:

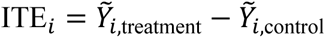

This framework effectively adjusts for variations in intrinsic cell features, providing a robust estimation of the ITE.

### CATE/CACRS estimation and statistical test

To calculate the CATE for a subgroup 𝑆 (e.g. cells of the same cell type), the average ITE across all cells in 𝑆 for gene *j* is computed as:

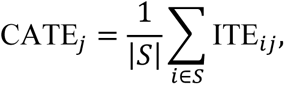

where |𝑆| denotes the number of cells in subgroup 𝑆.

To evaluate whether the conditional average treatment effect (CATE) for gene *j* is significantly greater than zero, we begin by formulating the null hypothesis that the observed individual treatment effect (ITE_*ij*_) values are equally likely to be positive or negative since the true value is 0 and there is continuous random noise. To test this hypothesis, each ITE_*ij*_ is transformed into a binary variable *Z*_*ij*_, where

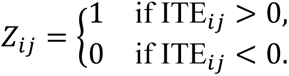

Under the null hypothesis, these binary variables follow a Bernoulli distribution with 𝑝 = 0.5. Next, the binary outcomes are aggregated within a subgroup 𝑆, where we calculate 𝑁_1_ by adding the count of positive outcomes (*Z*_*ij*_ = 1) with 0.5 times the count of *Z*_*ij*_ = 0. For large |𝑆|, the central limit theorem enables the approximation of 𝑁_1_by a normal distribution with mean 𝜇 = 0.5|𝑆| and variance 𝜎^2^ = 0.25|𝑆|. The z-score is then computed to quantify the deviation of 𝑁_1_ from its expected value, defined as

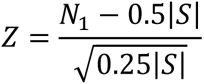

Finally, the p-value is calculated using the standard normal distribution’s cumulative distribution function,

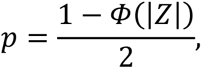

to assess the statistical significance of the observed deviation under the null hypothesis.

### Benchmarking datasets

#### PBMC dataset

This dataset comprises scRNA-seq data generated using the 10x Genomics platform, featuring multiplexed PBMC samples from SLE patients and controls^23^. Leveraging the demuxlet algorithm, it identifies cell origins and resolves doublets through genetic barcoding. The dataset consists of seven cell types, with 8,886 stimulated cells and 8,007 control cells, enabling comparisons between IFN-beta-stimulated and control conditions.

#### Rhinovirus infection dataset

Primary human bronchial epithelial cells were obtained from healthy adult donors and cultured at the air-liquid interface for 4 weeks to ensure maturation of cilia and mucus production^3^. The cells were then infected with mock or 1 × 10⁵ PFU human rhinovirus (HRV-01A) per organoid, with or without exposure to 2% cigarette-smoke extract (CSE). Single-cell suspensions were collected 5 days post-infection and processed using the 10x Genomics single-cell RNA-sequencing protocol. The dataset includes 26,420 cells across four conditions: mock (control condition), HRV, CSE, and HRV+CSE. We selected 7087 and 5168 cells under the CSE and mock condition for comparison and used the 10 annotated cell types provided by the authors for evaluation.

#### PBMC1 dataset^3^

Human PBMCs were isolated using Lymphoprep density gradient medium and plated at a density of 1 million cells per ml. The cells were stimulated for up to 48 hours with 1,000 U/ml of IFN-α2, IFN-β, IFN-γ, or IFN-III/IL-29, 100 ng/ml of IL-6, 20 ng/ml of TNF, or combinations of IFN-β with IL-6, TNF, or IFN-γ at the indicated concentrations for up to 48 h. We compared the 2759 and 2268 cells between the IFN-β condition and the ‘No stimulation’ condition from donor id ‘H3D2’. We used the 7 annotated cell types provided by the authors for evaluation.

#### ECCITE dataset

This dataset contains 20729 cells of 3 annotated cell cycle phases (G1, G2M, S) coming from 3 technique replicates measured using ECCITE-seq with 25 target genes generated from stimulated THP-1 cell line^9^. Since some previous methods only support the comparison between two conditions, we compared the 2386 negative control (NT) cells with the 18343 cells that are mutated for these methods. NT cells in 3 batches are treated as the control group. The annotated cell cycle phase was treated as cell type for model evaluation. We treated the 3 technique replicates as batches.

#### ASD dataset

This dataset was generated using in vivo Perturb-Seq to investigate the effects of 35 autism spectrum disorder (ASD) and neurodevelopmental delay (ND) risk genes on brain development^1^. CRISPR-Cas9-mediated frameshift mutations were introduced in these genes within the developing mouse neocortex in utero. Single-cell RNA sequencing was performed on 46,770 cortical cells at postnatal day 7 (P7) across 18 batches, focusing on five major cell types: cortical projection neurons, inhibitory neurons, astrocytes, oligodendrocytes, and microglia. We remove cells with annotated cell types other than the 5 cell types mentioned above since they are not perturbed, resulting in 40603 cells. For previous methods that only support the comparison between two conditions, we compared the 17813 negative control (annotated as ‘nan’) cells with the 22790 cells that are mutated for these methods. The ‘nan’ cells in the 18 batches are treated as the control group.

#### Type 1 diabetes dataset

We retrieved the scRNA-seq data of the T1D study from Fasolino et al.^49^. There are 69,645 cells from 24 batches under 3 conditions, where cultured pancreatic islet cells were sequenced from 11 healthy individuals, 5 individuals with T1D, and 8 individuals with no clinical presentation of T1D but positive for β-cell autoantibodies (AAB+). We use the “cell_type” annotation with 10 cell types provided by the authors for model evaluation. All cells from the Control sample are treated as the control group.

#### Human kidney multiomics atlas dataset

We retrieved the scRNA-seq and snRNA-seq data from the GEO under accession number GSE211785^50, 51^ (the PreSCVI version). This dataset contains 282,610 cells from 36 donors and 47 batches with 24 ‘Control’ batches and 23 ‘Disease’ batches, where 29 of them are scRNA-seq data and 18 of them are snRNA-seq data. Data were downloaded on December 15, 2024. All cells from the 24 ‘Control’ batches including both scRNA-seq and snRNA-seq data are treated as the control group. We used the “Cluster_Idents” annotation in the given data as the cell type annotation for model evaluation.

#### Mouse radiation therapy dataset^12^

This dataset comprises single-cell RNA sequencing data from the intestinal epithelium and lamina propria of C57BL/6J mice subjected to whole-abdominal irradiation or sham irradiation. Standard proton radiation therapy (0.9 ± 0.08 Gy s⁻¹) was delivered, and samples were collected on days 2, 3.5, 10, and 20 post-irradiation. Single cells from each condition were pooled, and flow cytometry was used to enrich 10,000 live cells per fraction without targeting specific populations. Single-cell libraries were prepared using the 10x Chromium platform and sequenced on an Illumina NextSeq. There are 91752 cells from the epithelial and lamina propria layers of the intestinal segments sequenced at 4 distinct times by two separate technical groups from mice under the control and radiation-treated condition. We use the “compartment” annotation as the cell type annotation for the data distortion analysis, and we combine the batch-related factors including the time at sequencing (annotation “day”), technical group (annotation “source”), and annotation “replicate” as the batch label and the annotation “sample” as the condition label. Data was downloaded on 13^th^ December 2024.

#### SEA-AD dataset^26^

We retrieved the “Whole Taxonomy - MTG: Seattle Alzheimer’s Disease Atlas (SEA-AD)” dataset from the cellxgene database. This dataset contains 1,378,211 single-nucleus RNA sequenced cells from the middle temporal gyrus (MTG) region from 47 donors marked as ’normal’ and 42 donors marked as ’dementia’ in the ’disease’ annotation. We use the “Subclass” annotation as the cell type annotation for downstream analysis and all cells from the 47 donors marked as ’normal’ are treated as the control group.

## Data availability

The PBMC dataset is available from the GEO under accession number GSE96583 and can be downloaded from the scGen tutorial https://drive.google.com/uc?id=1r87vhoLLq6PXAYdmyyd89zG90eJOFYLk. The rhinovirus infection dataset and another PBMC dataset are available at https://datadryad.org/stash/dataset/doi:10.5061/dryad.4xgxd25g1. The ECCITE-seq dataset is available from the GEO under accession number GSE153056 and can be downloaded from pertpy^2^ by the code “pertpy.data.papalexi_2021()”. The ASD dataset is available from the GEO under accession number GSE157977. The T1D dataset is available from the GEO under accession number GSE148073 and can be downloaded from https://cellxgene.cziscience.com/collections/51544e44-293b-4c2b-8c26-560678423380. The kidney dataset is available from the GEO under accession number GSE211785 and we use the version “GSE211785_Susztak_SC_SN_ATAC_merged_PreSCVI_final.h5ad.gz”. The mouse radiation therapy dataset is available from the GEO under accession number GSE280883. The SEA-AD MTG snRNA-seq dataset is available at https://cellxgene.cziscience.com/collections/1ca90a2d-2943-483d-b678-b809bf464c30.

## Code availability

The NDreamer package is available at https://github.com/lugia-xiao/NDreamer with detailed tutorials. All codes used to reproduce the results shown in the article is available at https://github.com/lugia-xiao/NDreamer_reproducible.

## Supplementary Tables

**Supplementary Table 1.** Original data for the quantitative evaluation in the experimental perturbation studies.

**Supplementary Table 2.** Original data for the quantitative evaluation in the batch effect removal part.

**Supplementary Table 3.** The top 300 up- and down-regulated genes with the smallest BH- adjusted p-values for each cell type in the SEA-AD dataset.

## Supporting information

https://drive.google.com/file/d/1I2kXyajm5ZdOBvLOQnHFBbN7PfBWf86b/view?usp=drive_link

https://drive.google.com/file/d/1BnLIElDTMNU7yU5MnNOKrsc3WC4botXM/view?usp=drive_link

---

**Extended Data Figure 1.**
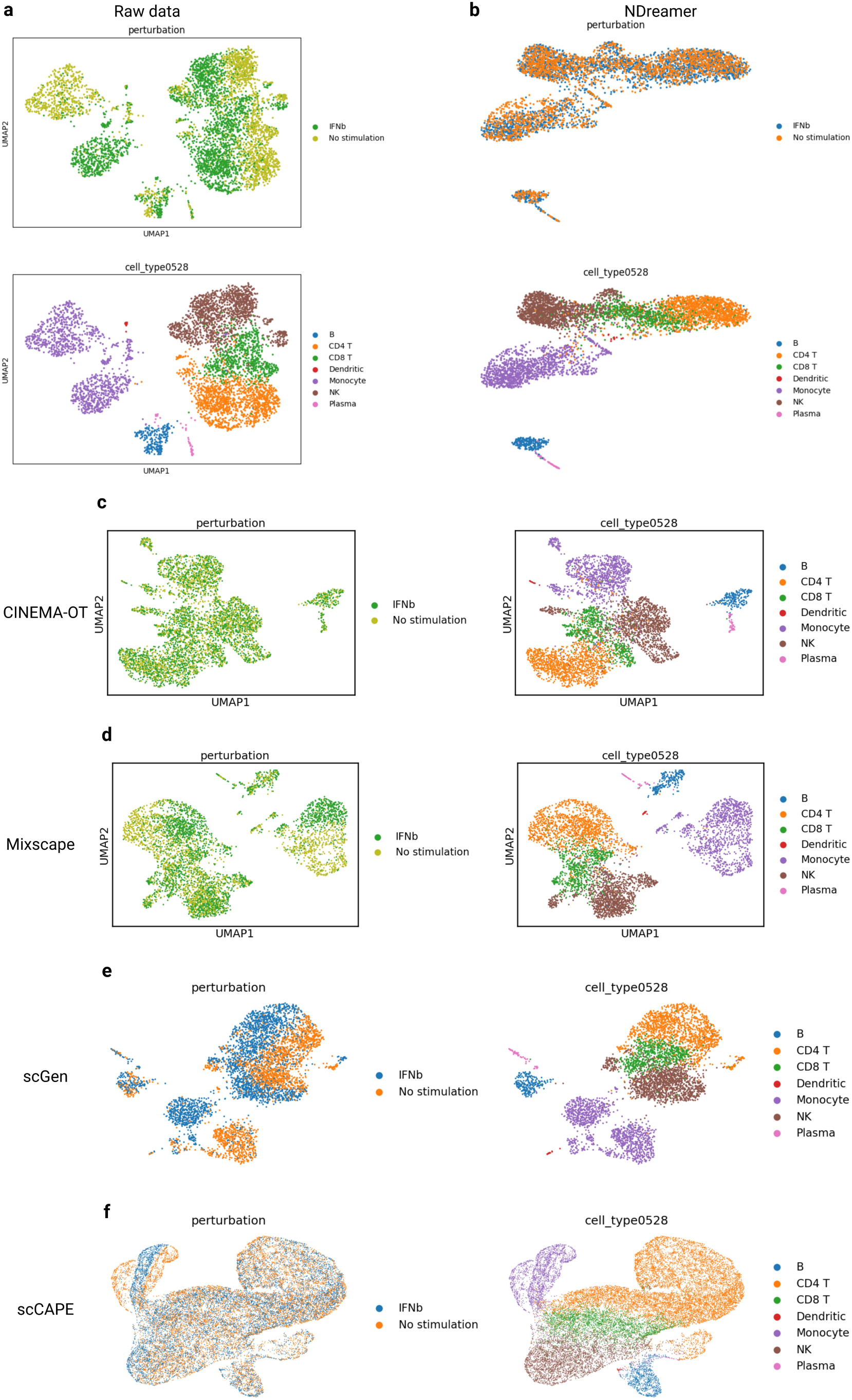
The UMAP visualization of cells in the PBMC1 dataset colored by cell type annotations and conditions for the **(a)** raw expression data, and the condition-free embeddings of **(b)** NDreamer, **(c)** CINEMA-OT, **(d)** Mixscape, **(e)** scGen, and **(f)** scCAPE.

**Extended Data Figure 2.**
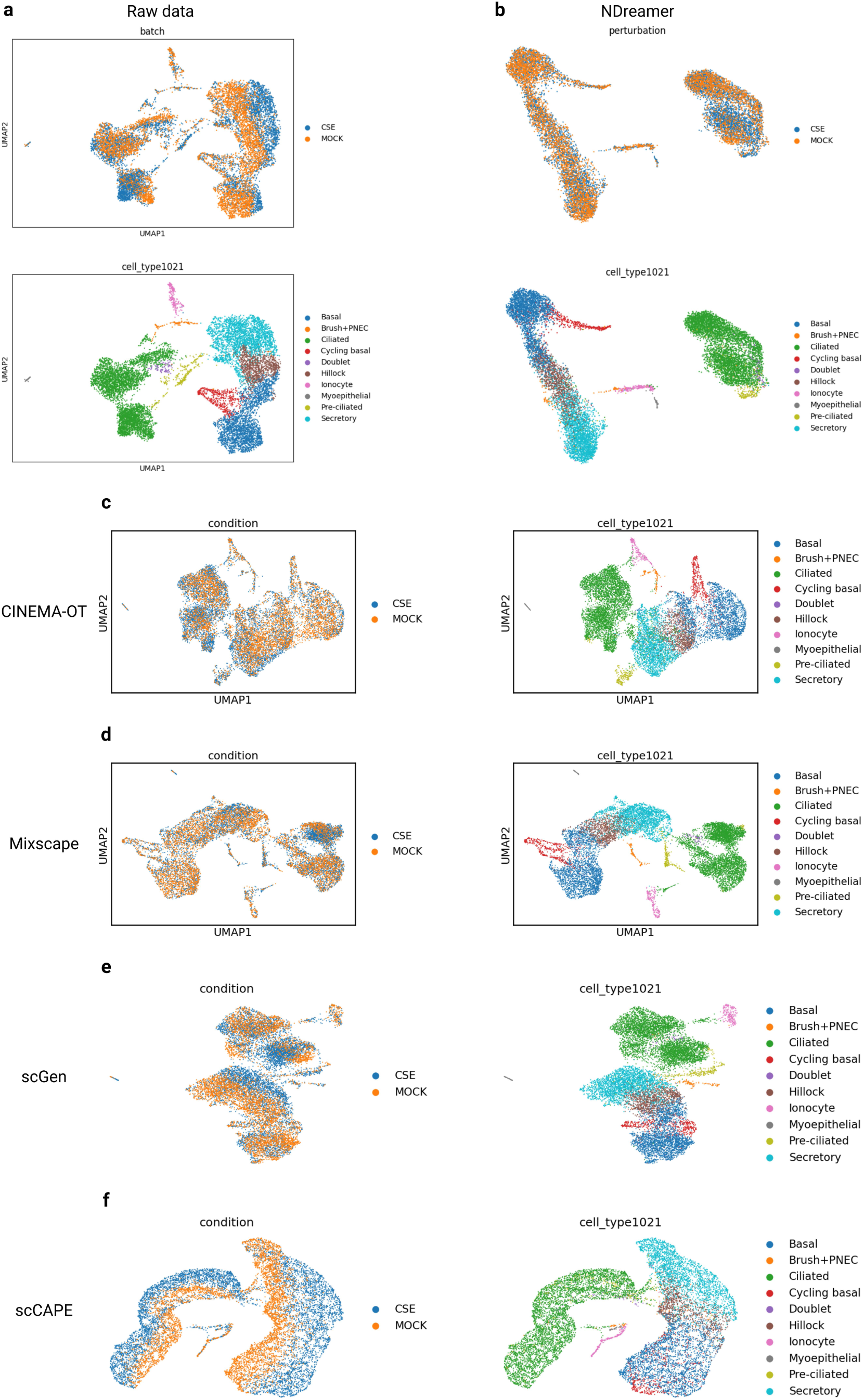
The UMAP visualization of cells in the virus dataset colored by cell type annotations and conditions for the **(a)** raw expression data, and the condition-free embeddings of **(b)** NDreamer, **(c)** CINEMA-OT, **(d)** Mixscape, **(e)** scGen, and **(f)** scCAPE.

**Extended Data Figure 3.**
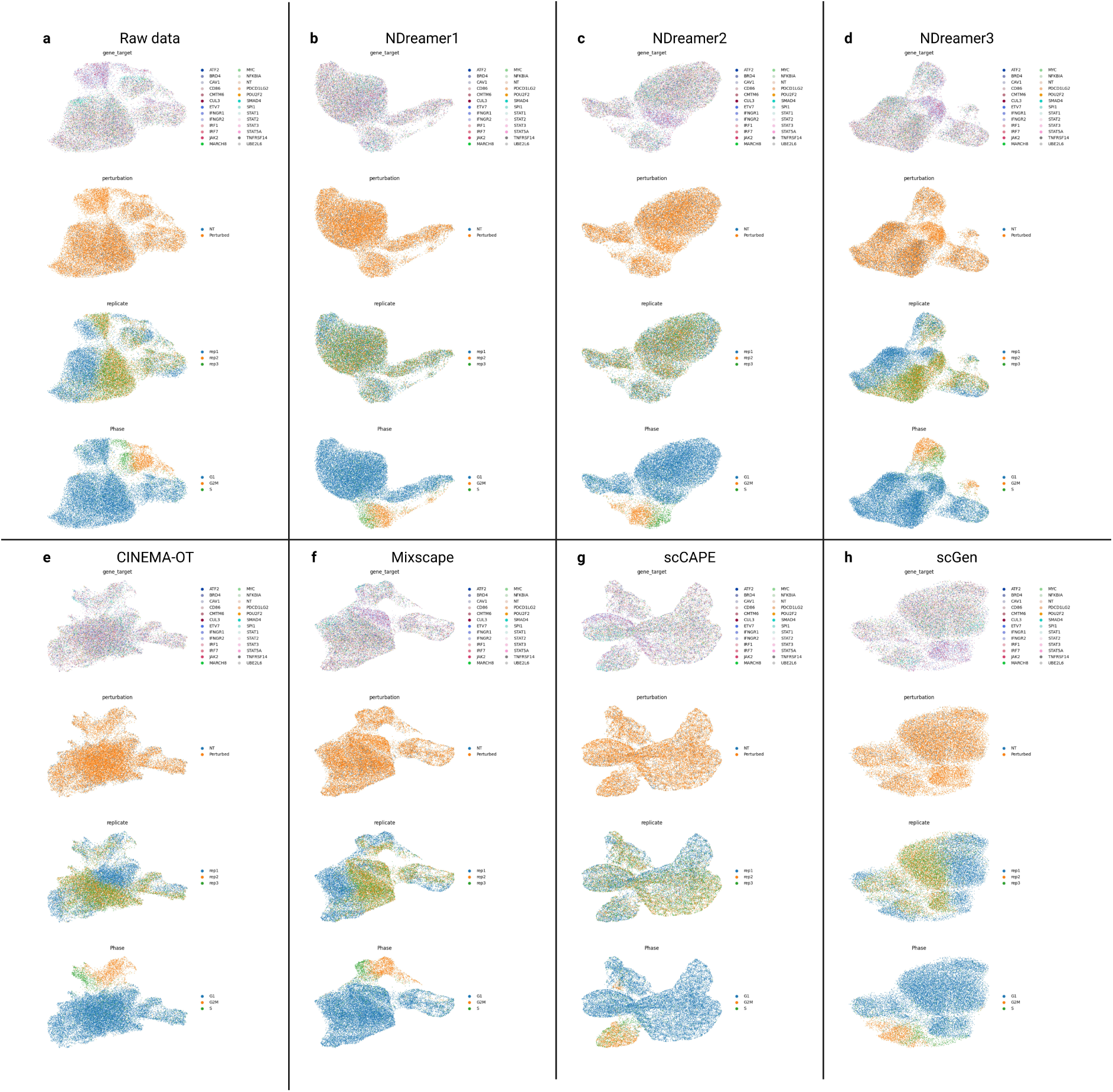
The UMAP visualization of cells in the ECCITE dataset colored by the target gene of the CRISPR, whether or not this cell is genetically perturbed, batch, and cell cycle phase annotations for the **(a)** raw expression data, and the condition-free embeddings of **(b)** NDreamer with batch effect removal and treating perturbed target genes as conditions (NDreamer1), **(c)** NDreamer with batch effect removal and treating perturbed target genes as conditions, which is the one we used in Figure 3 (NDreamer2), **(d)** NDreamer without batch effect removal and treating whether or not one cell is genetically perturbed as condition (NDreamer3), **(e)** CINEMA-OT, **(f)** Mixscape, **(g)** scCAPE, and **(h)** scGen.

**Extended Data Figure 4.**
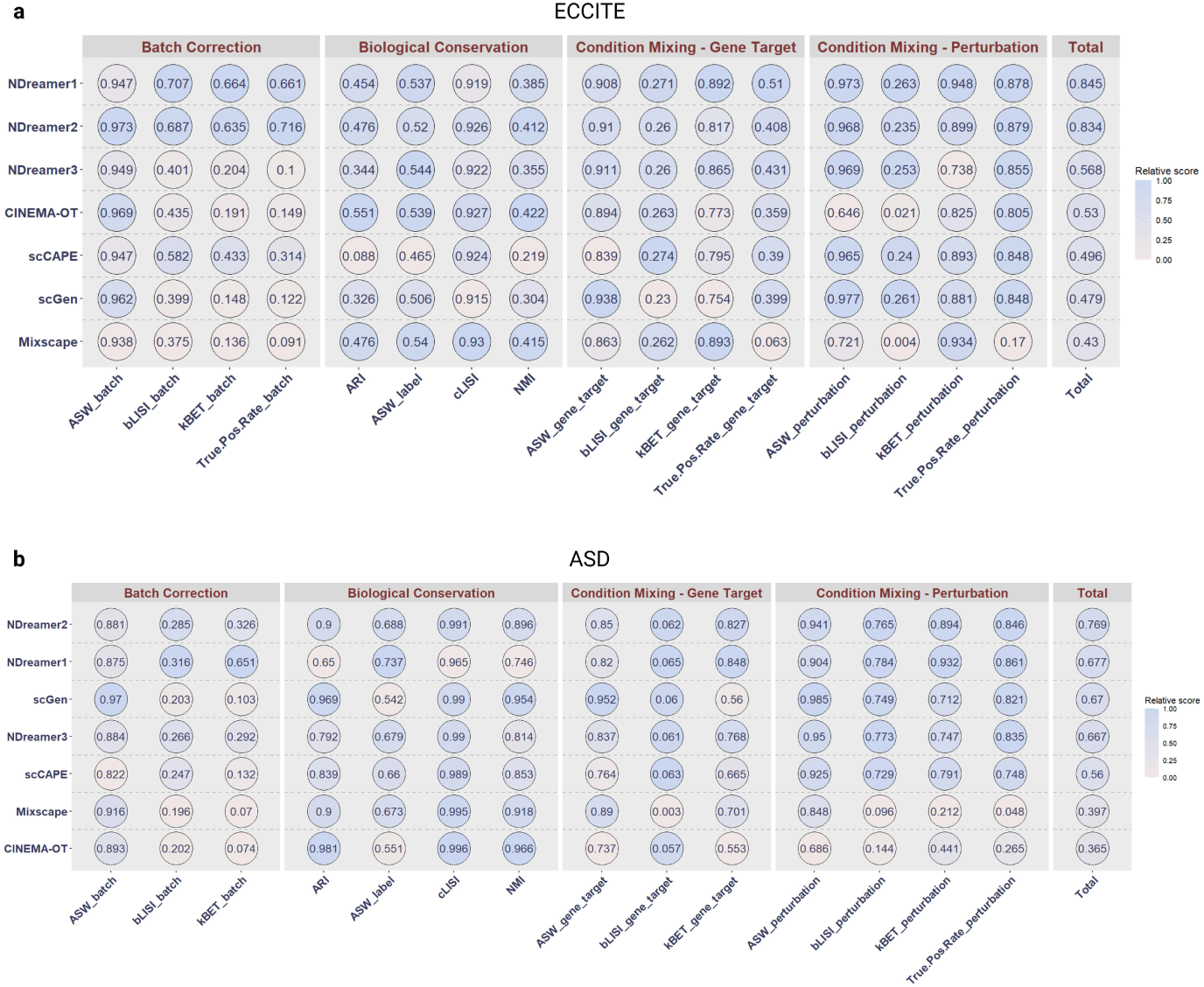
Quantitative benchmarking when treating each genetic mutation as a condition and enabling batch-effect-removal, if applicable. The evaluation metrics include biological conservation, batch-effect-removal (condition-mixing metrics applied to batch), condition-mixing metrics applied to perturbed CRISPR target genes, condition-mixing metrics applied to whether or not one cell is genetically perturbed, and their overall scores. The evaluations were performed for the **(a)** ECCITE and **(b)** ASD datasets.

**Extended Data Figure 5.**
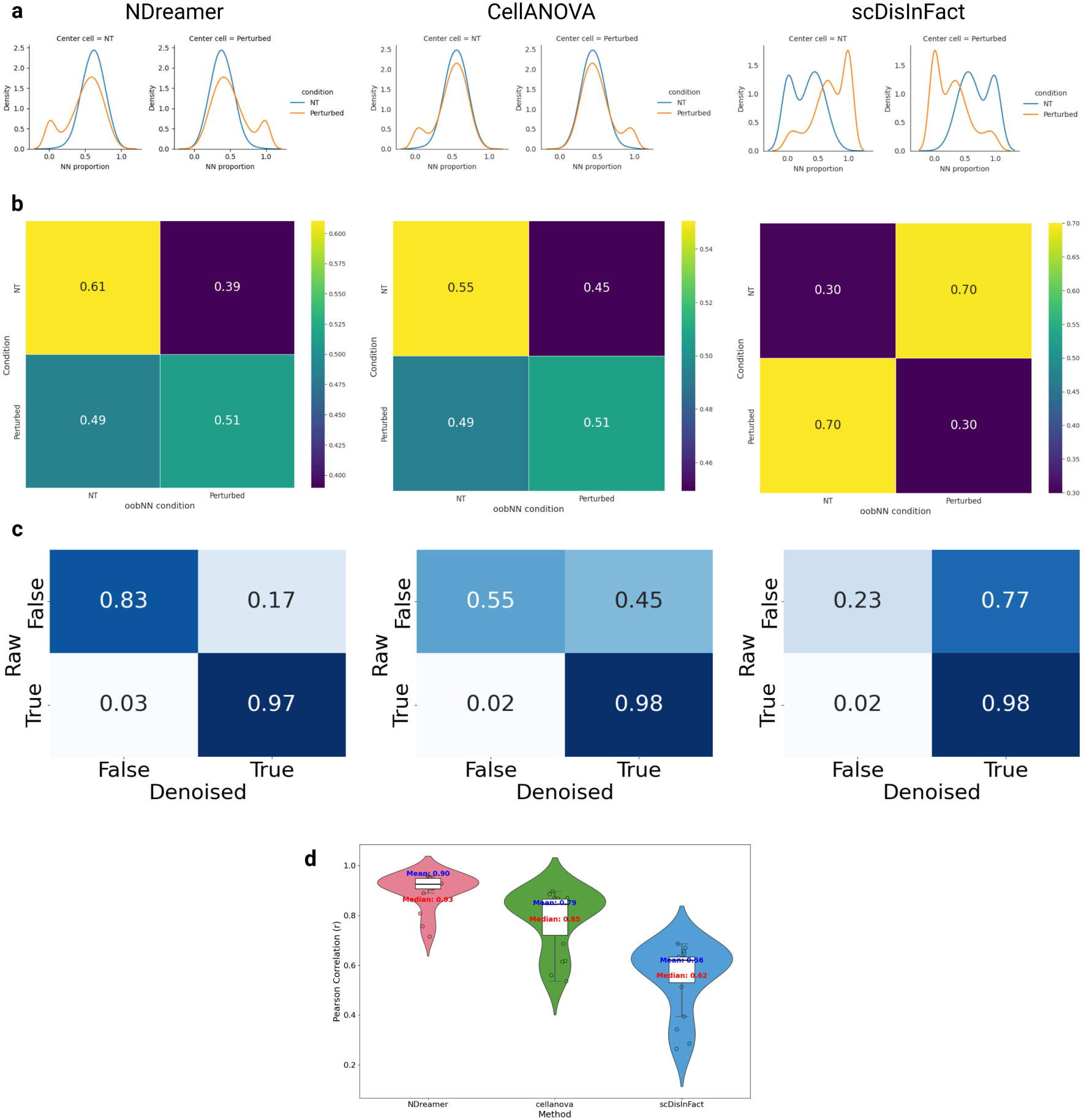
Evaluation of NDreamer, CellANOVA, and scDisInFact in producing batch-effect-free and condition-related-signal-preserved expression for the ECCITE dataset. **a.** The distribution of the oobNN condition proportion for each condition. **b.** The neighboring cell condition versus central cell condition matrix visualization. **c.** Visualization of the normalized confusion matrix between the DEGs with BH-adjusted p-values less than 0.05 in the raw and denoised expression. **d.** The distributions of the PCC between raw and denoised expression’s each gene’s logit(adjusted p-value) obtained from DEG analysis for each cell type in each CBC across 3 methods.

**Extended Data Figure 6.**
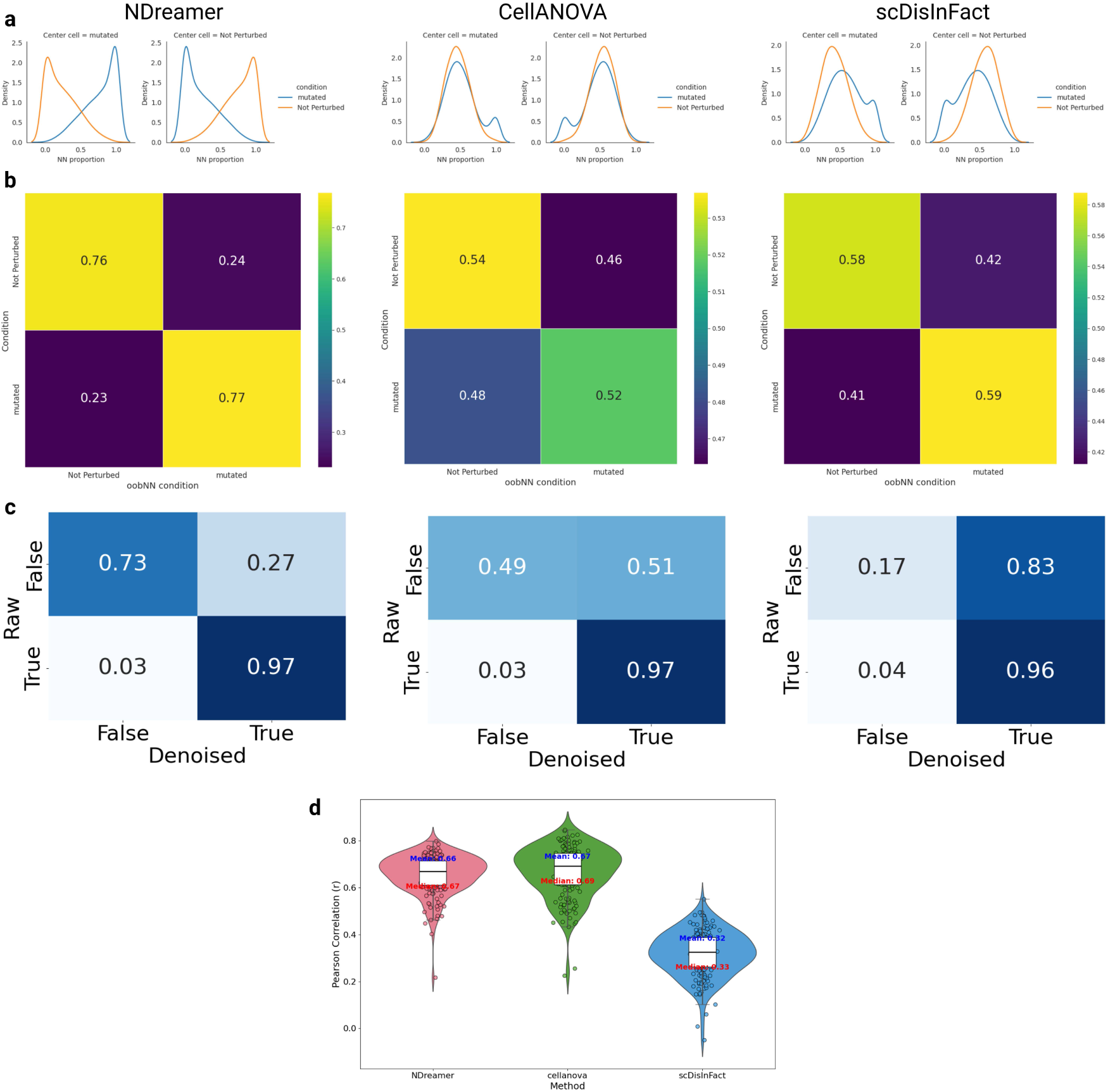
Evaluation of NDreamer, CellANOVA, and scDisInFact in producing batch-effect-free and condition-related-signal-preserved expression for the ASD dataset. **a.** The distribution of the oobNN condition proportion for each condition. **b.** The neighboring cell condition versus central cell condition matrix visualization. **c.** Visualization of the normalized confusion matrix between the DEGs with BH-adjusted p-values less than 0.05 in the raw and denoised expression. **d.** The distributions of the PCC between raw and denoised expression’s each gene’s logit(adjusted p-value) obtained from DEG analysis for each cell type in each CBC across 3 methods.

**Extended Data Figure 7.**
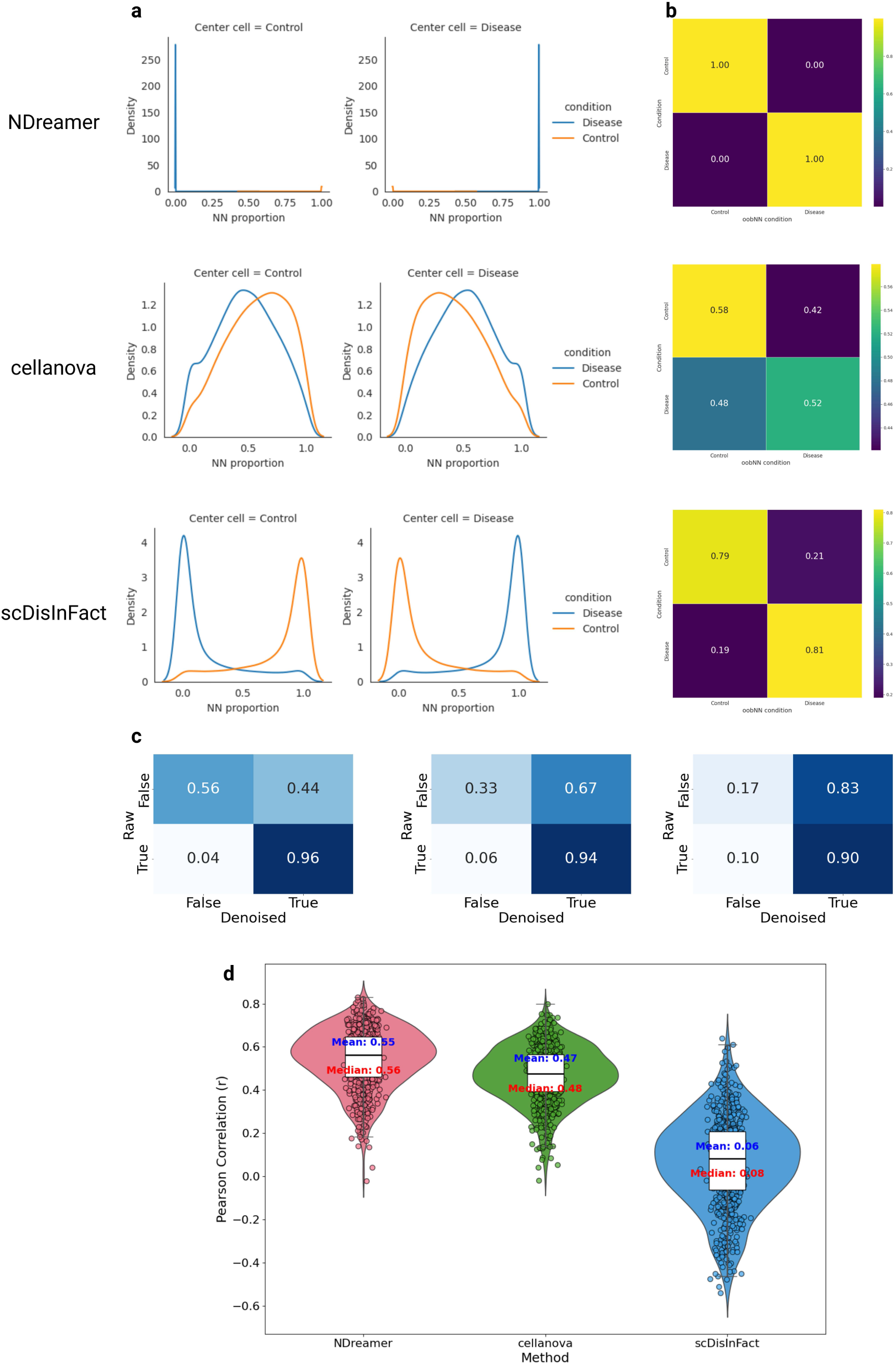
Evaluation of NDreamer, CellANOVA, and scDisInFact in producing batch-effect-free and condition-related-signal-preserved expression for the kidney dataset. **a.** The distribution of the oobNN condition proportion for each condition. **b.** The neighboring cell condition versus central cell condition matrix visualization. **c.** Visualization of the normalized confusion matrix between the DEGs with BH-adjusted p-values less than 0.05 in the raw and denoised expression. **d.** The distributions of the PCC between raw and denoised expression’s each gene’s logit(adjusted p-value) obtained from DEG analysis for each cell type in each CBC across 3 methods.

**Extended Data Figure 8.**
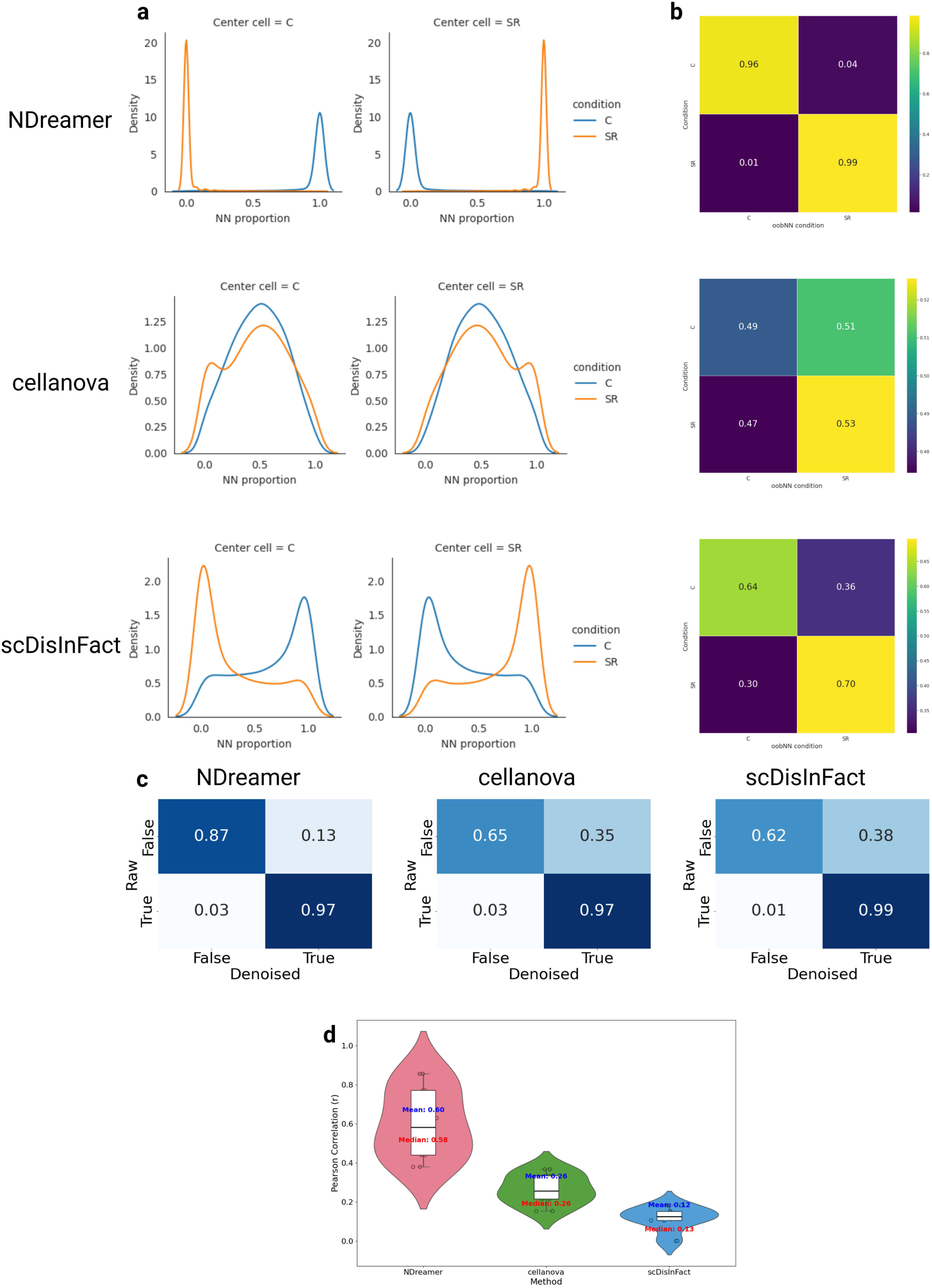
Evaluation of NDreamer, CellANOVA, and scDisInFact in producing batch-effect-free and condition-related-signal-preserved expression for the Mouse dataset. **a.** The distribution of the oobNN condition proportion for each condition. **b.** The neighboring cell condition versus central cell condition matrix visualization. **c.** Visualization of the normalized confusion matrix between the DEGs with BH-adjusted p-values less than 0.05 in the raw and denoised expression. **d.** The distributions of the PCC between raw and denoised expression’s each gene’s logit(adjusted p-value) obtained from DEG analysis for each cell type in each CBC across 3 methods.

**Extended Data Figure 9.**
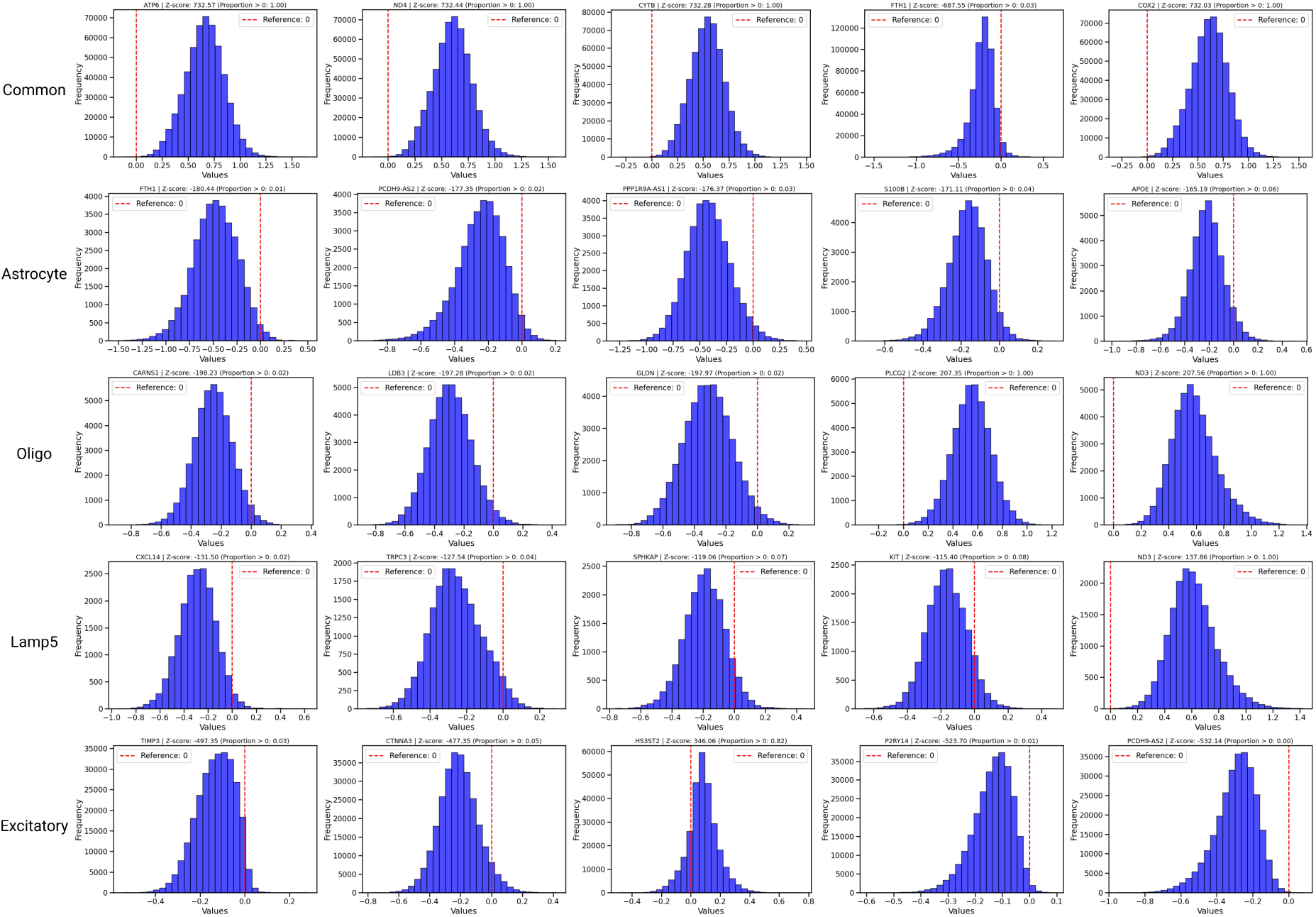
Visualization of the CRS distribution for genes and cell types mentioned in the SEA-AD dataset.

**Extended Data Figure 10.**
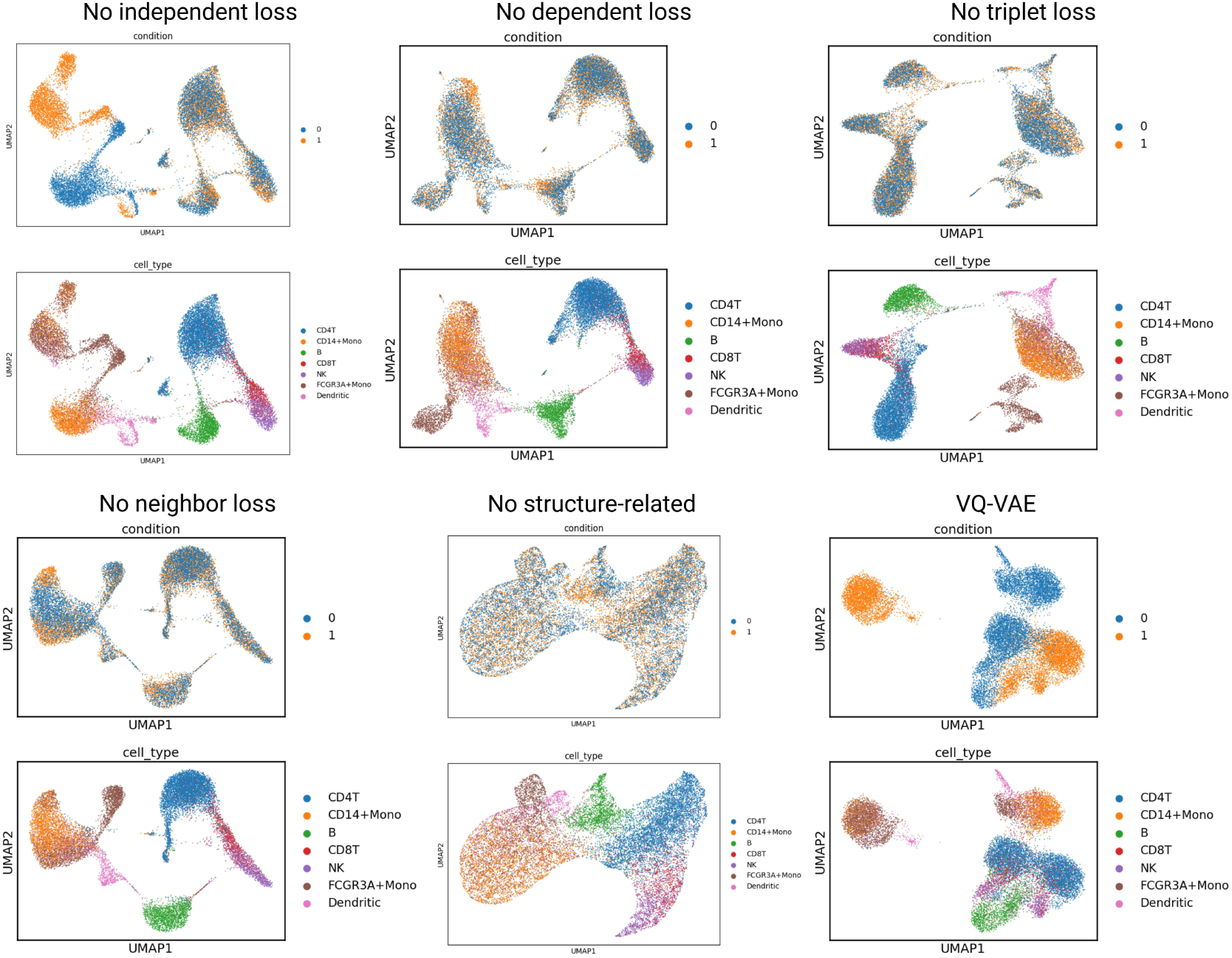
UMAP visualization of the models mentioned in the ablation study for the PBMC dataset colored by cell type annotations and conditions.

