## Supplementary material for "Single-cell-level condition-related signal estimation with batch effect removal through neural discrete representation learning": https://drive.google.com/file/d/1I2kXyajm5ZdOBvLOQnHFBbN7PfBWf86b/view?usp=drive_link

### Supplementary Notes

Xiao Xiao

January 2025

### 1 List of Notations and default hyperparameter settings regarding the model structure

#### 1.1 Model input

Here we talk about the situation of a single cell input.

- Cell $i$ have its expression, represented by a vector $Y_i \in \mathbb{R}^{n_{hvg}}$, batch $b_i$ (scaler), and condition $c_i$ (scaler).
- $n_{hvg}$ is the number of selected highly variable genes, default to 2000.

#### 1.2 Encoder

Here we talk about the situation of a single cell input.

- In our model, there are $n_c$ codebooks, each contains $n_e$ embeddings $e$ of dimensionality $d_c$.
- $n_c$ is default to 32, $n_e$ is default to 1024, $d_c$ is default to 8. So there are almost $1024^{32}$ of possible combinations, which is much more than the number of cells of all humans on earth.
- Within one codebook, $Y_i$ is transformed by a codebook-specific neural network into an intermediate latent space $Z_i \in \mathbb{R}^{d_c}$.
- All codebook-specific neural networks have the input dim of $n_{hvg}$ and output dim of $d_c$. They shared the parameters in the first 2 hidden layers with default values of [1024, 512]. The RELU activation function and BatchNorm1D are used between each layers.
- Putting all codebooks together, we get the codebook selection recording matrix $P$ of shape $n_c \times n_e$. We then concatenate the chosen embeddings from each codebook to form a new resampled latent space $Z_i' \in \mathbb{R}^{n_c d_c}$.
- In the commitment loss, $\text{sg}(\cdot)$ is the stop-gradient operator and $\beta$ (default to 0.25) controls the balance between the two terms.

- In the variational inference part, the mean vector $\mu_i \in \mathbb{R}^{d_z}$ and the variance vector $\sigma_i \in \mathbb{R}^{d_z}$, where $d_z$ refers to the number of dimensions of the effect modifier embedding. Here, $\mu_i$ is the effect modifier embedding $X_i$.
- $d_z$ is default to 256.
- In the re-sampling part, $\epsilon$ refers to the random gaussian noise.

##### 1.3 Decoder

Here we talk about the situation of a single cell input.

- $E_{b_i}, E_{c_i}$ refers to the embedding of batch and condition for cell $i$, their default embedding dim is 256.
- $\mathrm{f}_{\mathrm{recon}}$ refers to the decoder neural network. With the input dimension of $d_z$ and the output dimension of $n_{hvg}$, the default number of hidden layers is [512,1024], with RELU activation function between each hidden layer.
- $\hat{Y}_i$ refers to the reconstructed expression.

#### 2 Hyperparameter settings

The hyperparameter settings are the same across all datasets mentioned in the article. The running times reported in the discussion part of the article were based on the A100 GPU. The running time increases linearly with the number of cells, number of batches, and number of conditions.

Table 1: Default Values of Hyperparameters

| **Hyperparameter** | **Default Value** |
|---|---|
| $\lambda_{\mathrm{commit}}$ | 1 |
| $\lambda_{\mathrm{KL}}$ | 0.005 |
| $\lambda_{\mathrm{recon}}$ | 1 |
| $\lambda_{\mathrm{cos}}$ | 20 |
| $\lambda_{\mathrm{dependent}}$ | 50 |
| $\lambda_{\mathrm{independent}}$ | 1000 |
| $\lambda_{\mathrm{triplet}}$ | 5 |
| $\lambda_{\mathrm{neighbor}}$ | 1 |
| $k_{\mathrm{neighbor}}$ | 20 |
| Batch size | 1024 |
| Epoch | 15 |

### 3 Advantages of neural discrete representation learning

- In a continuous embedding, it is hard to establish independence between the effect modifier embedding and the batch (or condition). We may be able to make the parametric or non-parametric correlation coefficient between a set of embeddings and batch very low. But we can always find another transformation that switches the correlation relationship between the embedding and the condition. And it is unclear that whether a 0 or low correlation coefficient would really indicate independence (e.g. y=sin(x), $-2\pi \leq x \leq 2\pi$). However, for discrete latent variables, we can directly measure the independence by calculating the mutual information between two probability matrices, where 0 mutual information directly indicates independence.
- It is also hard to say if the embedding generated by models with discriminator is really independent of condition or batch. When trying to use a discriminator to establish independence, it is unclear whether the discriminator would really extract all information in the embedding that is related to the batch or condition. This is determined by the hyperparameter settings of the discriminator (e.g. the ratio of the learning rate between the generator and the discriminator and the training schedule).
- Adversarial training takes a lot of time since it requires multiple times of backward propagation, while the loss function of mutual information does not cost much time.

### 4 Proof of bias in weighting-based ITE estimation

Consider the experimental perturbation scenario where there is no significant difference in the expression of gene $x$ between the case and control groups. The library-size normalized expression is continuous and non-negative. Given the sparsity of single-cell sequencing data, there is a probability that this gene is not detected (dropout). Without loss of generality, the expression $x_i$ of this gene in a single cell $i$ follows the distribution:

$$p(x_i) = \begin{cases} 0, & x_i < 0 \\ p, & x_i = 0 \\ f(x_i), & x_i > 0 \end{cases}$$

where $0 < p < 1$ and $f(x_i)$ represents an arbitrary density function such that

$$\int_0^\infty f(x_i)\, dx_i = 1 - p.$$

Since there is no significant difference in the expression of gene $x$ between the case and control groups, we suppose that the expressions of gene $x$ in all cells, no matter in the case or control group, are independent and identically distributed (i.i.d).

In weighting-based causal inference methods, let $k_i \in \mathbb{N}$ cells, indexed as $c_1, c_2, \ldots, c_{k_i}$, be used to compute $x_i'$, the counterfactual expression of gene $x$ in cell $i$. The corresponding weights assigned to these $k_i$ cells are $w_1, w_2, \ldots, w_{k_i}$ and satisfy the constraint $\sum_{j=1}^{k_i} w_j = 1$. The individual treatment effect (ITE) is then estimated as:

$$\hat{\text{ITE}} = x_i - x_i' = x_i - \sum_{j=1}^{k_i} w_j x_{c_j}.$$

We now analyze the probability of different expression scenarios:

- The probability that cell $i$ has nonzero expression while its counterfactual cell has zero expression:

$$P(x_i > 0, x_i' = 0) = (1-p)p^{k_i}.$$

- The probability that both cell $i$ and its counterfactual cell have nonzero expression:

$$P(x_i > 0, x_i' > 0) = (1-p)(1-p^{k_i}).$$

- The probability that both cell $i$ and its counterfactual cell have zero expression:

$$P(x_i = 0, x_i' = 0) = p^{k_i+1}.$$

- The probability that cell $i$ has zero expression while its counterfactual cell has nonzero expression:

$$P(x_i = 0, x_i' > 0) = p(1-p^{k_i}).$$

Next, we compute the probability that the estimated ITE is negative or exactly zero (with a weighting factor of $\frac{1}{2}$ for the zero cases as used in paired nonparametric test):

$$\begin{aligned}
&P(\hat{\text{ITE}} < 0) + \frac{1}{2}P(\hat{\text{ITE}} = 0) \\
&= P(x_i = 0, x_i' > 0) + \frac{1}{2}P(x_i = 0, x_i' = 0) + \frac{1}{2}P(x_i > 0, x_i' > 0) \\
&= p(1-p^{k_i}) + \frac{1}{2}p^{k_i+1} + \frac{1}{2}(1-p)(1-p^{k_i}) \\
&= \frac{1}{2} + \frac{p}{2} - \frac{p^{k_i}}{2}.
\end{aligned}$$

The inclusion of the term $\frac{1}{2}P(x_i > 0, x_i' > 0)$ arises due to the symmetry and the i.i.d. assumption of the distribution of $x$ across all cells.

From this result, we observe:

- For nearest neighbor matching, $k_i = 1$, the probability simplifies to

$$P(\hat{\text{ITE}} < 0) + \frac{1}{2}P(\hat{\text{ITE}} = 0) = 0.5,$$

indicating an unbiased estimate.

- For weighting-based methods, $k_i \neq 1$,

$$P(\hat{\text{ITE}} < 0) + \frac{1}{2}P(\hat{\text{ITE}} = 0) > 0.5,$$

demonstrating a systematic bias towards negative estimates.

Thus, weighting-based methods introduce bias in ITE estimation when $k_i \neq 1$.
