## Supplementary material for "Single-cell-level condition-related signal estimation with batch effect removal through neural discrete representation learning": https://drive.google.com/file/d/1BnLIElDTMNU7yU5MnNOKrsc3WC4botXM/view?usp=drive_link

### Supplementary Figures

Xiao Xiao

January 2025

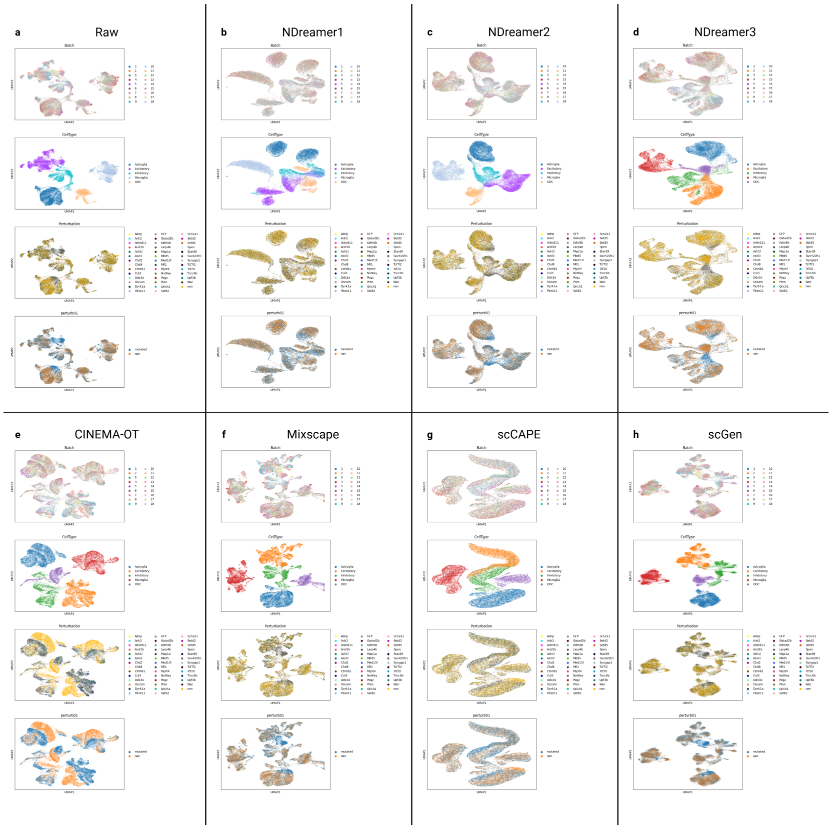


**Supplementary Figure 1:** The UMAP visualization of cells in the ASD dataset colored by batch, cell type, the target gene of the CRISPR, and whether or not this cell is genetically perturbed annotations for the **(a)** raw expression data, and the condition-free embeddings of **(b)** NDreamer with batch effect removal and treating perturbed target genes as conditions (NDreamer1), **(c)** NDreamer with batch effect removal and treating perturbed target genes as conditions, which is the one we used in Figure 3 (NDreamer2), **(d)** NDreamer without batch effect removal and treating whether or not one cell is genetically perturbed as condition (NDreamer3), **(e)** CINEMA-OT, **(f)** Mixscape, **(g)** scCAPE, and **(h)** scGen.

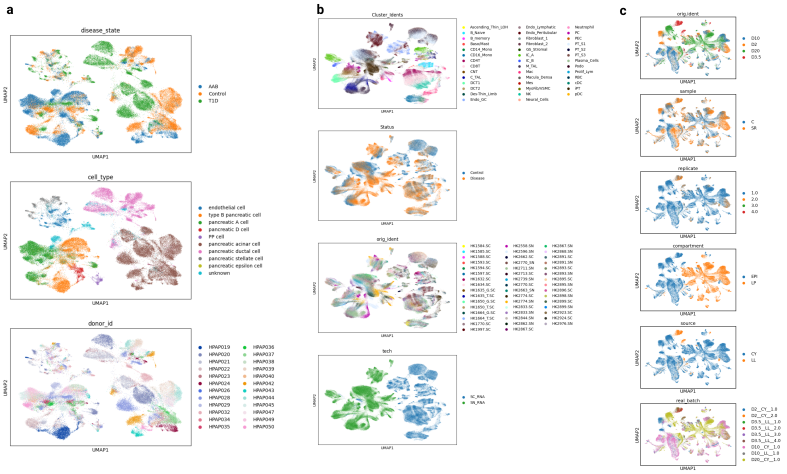


**Supplementary Figure 2: UMAP visualization of the raw expression for the T1D, kidney, and mouse dataset. a.** The UMAP visualization of the T1D dataset colored by disease state (condition), cell type, and batches. **b.** The UMAP visualization of the kidney dataset colored by cell type, condition, batch, and data modality. **c.** The UMAP visualization of the mouse dataset colored by the time of sequencing (orig.ident), sample (condition), replicate, compartment (raw cell type annotation), source (technical group), and the batch annotations.

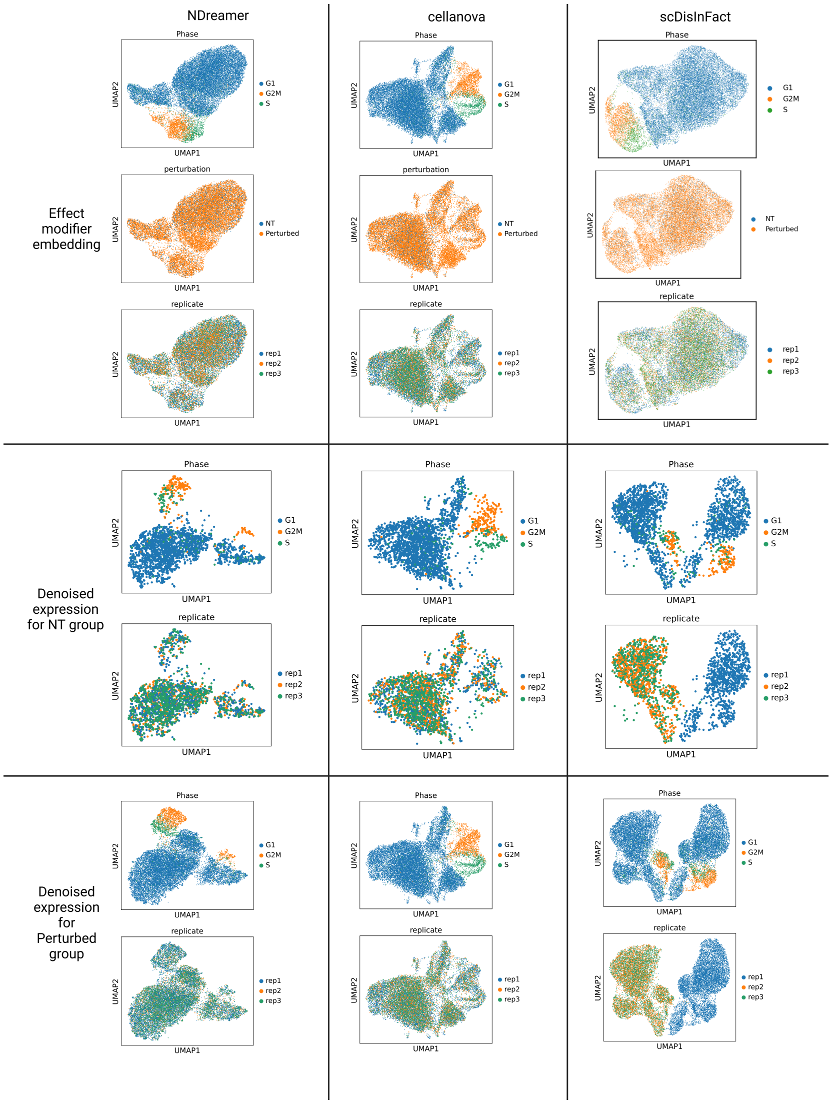


**Supplementary Figure 3:** UMAP visualization of the effect modifier embeddings, the denoised expressions for the negative control (NT) group, and the denoised expressions for the perturbed group (colored by batch, cell cycle phase, and condition annotation) for NDreamer, CellANOVA, and scDisInFact on the ECCITE dataset.

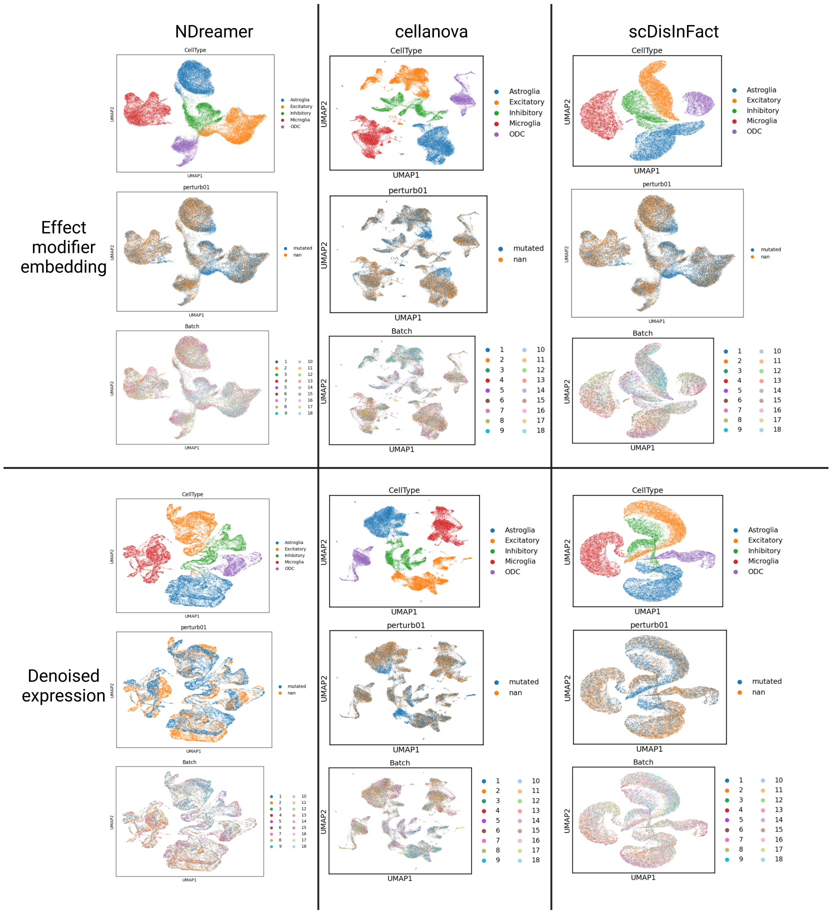


**Supplementary Figure 4:** UMAP visualization of the effect modifier embeddings and the denoised expressions colored by batch, cell type, and condition annotation for NDreamer, CellANOVA, and scDisInFact on the ASD dataset.

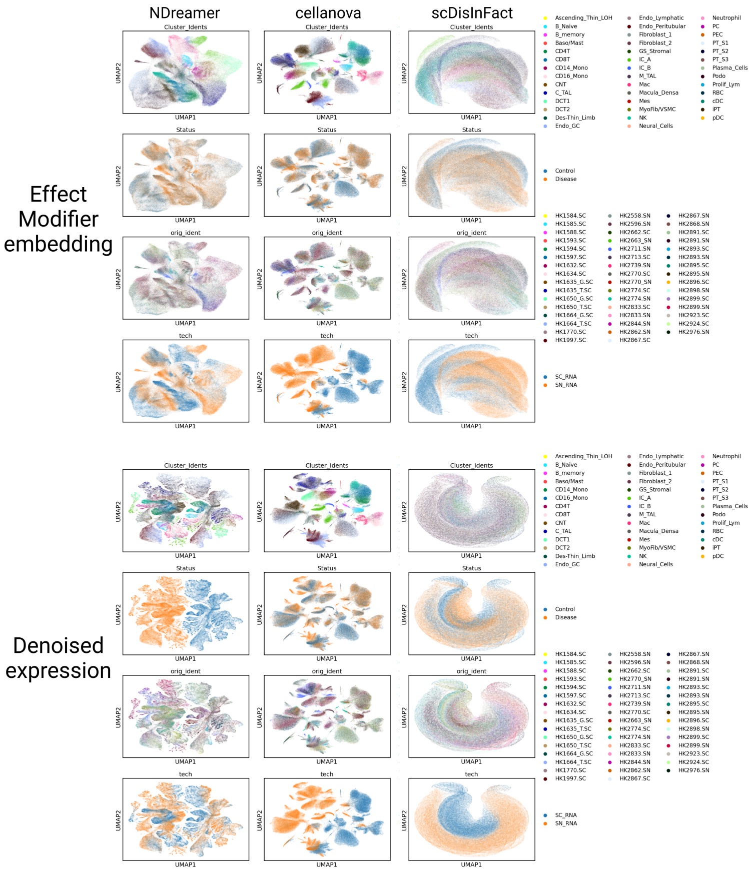


**Supplementary Figure 5:** UMAP visualization of the effect modifier embeddings and the denoised expressions colored by cell type, condition, batch, and data modality annotations for NDreamer, CellANOVA, and scDisInFact on the kidney dataset.

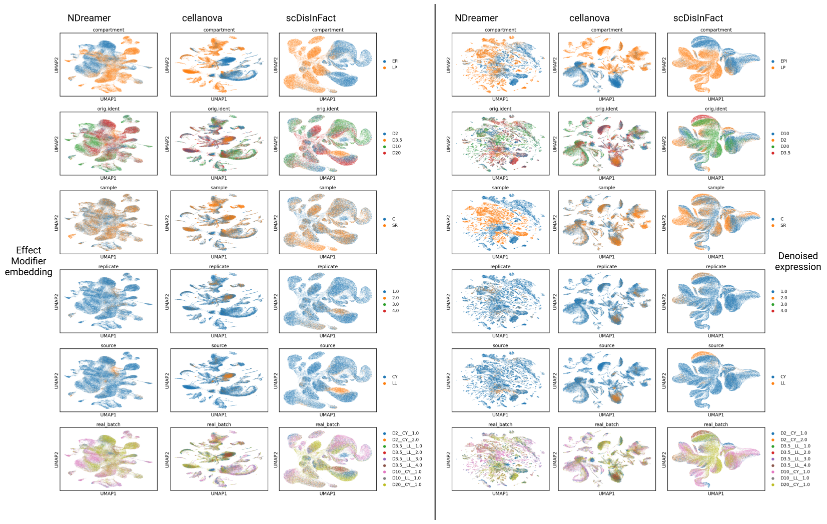


**Supplementary Figure 6:** UMAP visualization of the effect modifier embeddings and the denoised expressions colored by the time of sequencing (orig.ident), sample (condition), replicate, compartment (raw cell type annotation), source (technical group), and the batch annotations for NDreamer, CellANOVA, and scDisInFact on the mouse dataset.

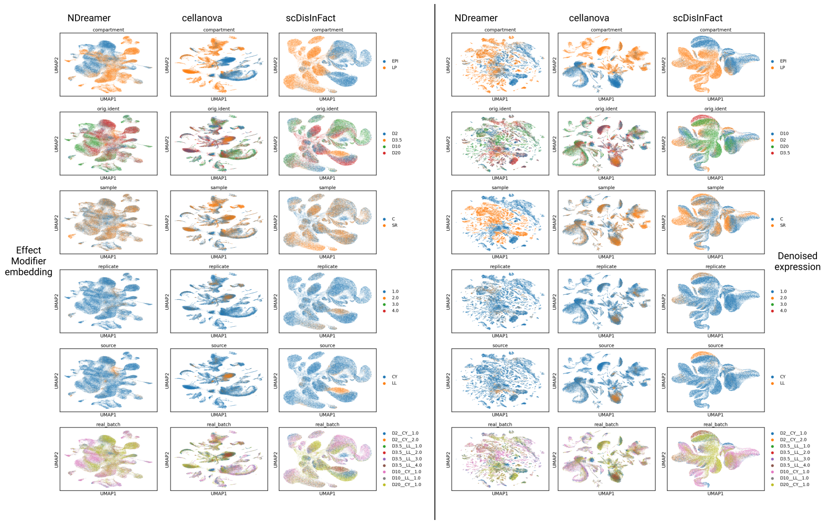


**Supplementary Figure 6:** UMAP visualization of the effect modifier embeddings and the denoised expressions colored by the time of sequencing (orig.ident), sample (condition), replicate, compartment (raw cell type annotation), source (technical group), and the batch annotations for NDreamer, CellANOVA, and scDisInFact on the mouse dataset.

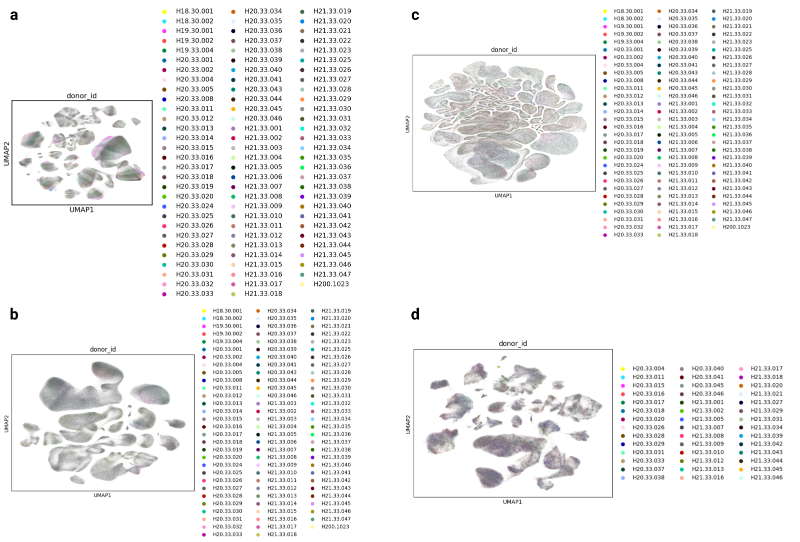


**Supplementary Figure 7:** UMAP visualization of the (a) raw expression, (b) effect modifier embeddings, (c) denoised expression, and (d) ITE colored by batch (donor_id)

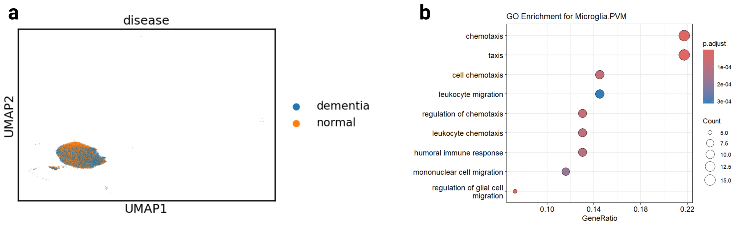


**c**

```
['CCL8', 'VCAN-AS1', 'CCL3', 'IFITM1', 'CH25H', 'SIGLEC1', 'H2BC7', 'FOS', 'RGS5', 'CYSLTR1', 'GASK1B-AS1', 'SELENOP', 'LINC027
12', 'LILRB5', 'JAML', 'H2BC8', 'VAV3', 'MRC1', 'HHATL', 'VCAN', 'NTN1', 'ZFP36L1', 'GPR183', 'FAM177B', 'LINC02642', 'LYZ', 'S
AT1', 'A2M', 'MNDA', 'MS4A4E', 'H1-3', 'CX3CR1', 'FTH1', 'PRKCQ', 'LPAR6', 'LTC4S', 'CD83', 'P2RY13', 'RNASE1', 'LPAR1', 'EPCAM
-DT', 'ELMO1-AS1', 'CA2', 'HSPA1A', 'PDK4', 'COLEC12', 'LINC00639', 'IER3', 'LOC285626', 'H2BC11', 'DUSP1', 'ZFP36', 'LINC0027
8', 'AIF1', 'CIMAP1B', 'PLPP3', 'ENO4', 'LINC00996', 'SHOC1', 'STON2', 'P2RY12', 'LINC01482', 'LINC01374', 'LOC124903386', 'CD2
2', 'ROBO2', 'IFI44L', 'SEMA6A', 'RMST', 'CSF2RA', 'SH3BP4', 'MROCKI', 'OLR1', 'LMCD1-AS1', 'TENT5A', 'VIM', 'HLA-DRB1', 'LINC0
1091', 'H2AC19', 'CCDC200', 'CCK', 'PLTP', 'XAF1', 'MIR646HG', 'ATF3', 'GNG12-AS1', 'LOC105373170', 'ATP8B4', 'H2BC4', 'HCG22',
'FOLR2', 'BTG2', 'SRGN', 'LINC02232', 'ITGA6', 'LINC01375', 'LINC02057', 'SAMD9L', 'EPAS1', 'NCKAP5']
```

**Supplementary Figure 8: ITE and CATE analysis for the microglia cell type in the SEA-AD dataset a.** UMAP visualization of the microglia cells colored by condition, with no significant differences between cells in two conditions. **b.** The enriched GO terms for the top 100 down-regulated genes for microglia. **c.** The list of the top 100 down-regulated genes.
